# Time-dependent Interaction Modification Generated from Plant-soil Feedback

**DOI:** 10.1101/2023.11.03.565336

**Authors:** Heng-Xing Zou, Xinyi Yan, Volker H.W. Rudolf

## Abstract

Pairwise interactions between species can be modified by other community members, leading to emergent dynamics contingent on community composition. Despite the prevalence of such higher-order interactions, little is known about how they are linked to the timing and order of species’ arrival. We generate population dynamics from a mechanistic plant-soil feedback model, then apply a general theoretical framework to show that the modification of a pairwise interaction by a third plant depends on its germination phenology. These time-dependent interaction modifications emerge from concurrent changes in plant and microbe populations and are strengthened by higher overlap between plants’ associated microbiomes. The interaction between this overlap and the specificity of microbiomes further determines plant coexistence. Our framework is widely applicable to mechanisms in other systems from which similar time-dependent interaction modifications can emerge, highlighting the need to integrate temporal shifts of species interactions to predict the emergent dynamics of natural communities.

## Introduction

Complex communities in nature often reassemble every growing season. The relative timing of species life history events in each season, i.e., phenology, can affect species interactions and community structure (Fukami 2015; Rudolf 2019; Zou & Rudolf 2023a). In pairwise systems, the shift in species interactions and community states with changes in species arrival times is well described by the framework of priority effects. However, theories studying the effects of variation in arrival sequence on pairwise systems (Song *et al*. 2020; Zou & Rudolf 2023b) have fallen behind the numerous experiments conducted in multispecies communities (Dickie *et al*. 2012; Drake 1991; Pu & Jiang 2015; Song *et al*. 2021b; Uricchio *et al*. 2019). The focus on pairwise interactions limits our perspective on widespread higher-order interactions that arise when the presence of a third species affects the interactions between two species (Billick & Case 1994; Werner & Peacor 2003; Wootton 1994) and are important to maintaining biodiversity (Levine *et al*. 2017). As species-rich communities in nature face widespread reshuffling in phenology and timing of interactions (Kharouba *et al*. 2018; Parmesan 2006), we need a general framework for higher-order interactions that includes this temporal dimension of community structure.

Documented in systems from microbes and plants to crustaceans and amphibians (Blackford *et al*. 2020; Fragata *et al*. 2022; Grainger *et al*. 2019; Rudolf 2018; Shorrocks & Bingley 1994), priority effects are generated by either positive frequency dependence or temporal changes in species traits due to different arrival times. Here, “arrival” can refer to both colonizations over many generations such as succession, and recurring arrival time such as the annual reassembly of communities (Zou & Rudolf 2023a). For example, early-colonizing bacteria may grow to higher abundance and exclude late arrivers (Grainger *et al*. 2019), and plants can be affected by the chemical composition or microbiomes in soil previously occupied by other species (Kardol *et al*. 2007). Because these changes in relative abundances or traits are not instantaneous but develop over time, the relative arrival time between two species directly modifies their interaction strengths. Similarly, in three-species communities, how the third species affects a pairwise interaction (interaction modification) can depend on its arrival time, which can change its density, traits, or both, and that of other community members. For instance, increasing the coverage of barnacles decreases bird predation of limpets in intertidal communities (Wootton 1993). If the coverage (*density*) of barnacles depends on the length they colonized the patch before limpets arrive, then this change in coverage over time could lead to a temporal change in the strength of interaction modification. In an aquatic predator-prey system, morphological changes of snails induced by the presence of a first predator increased its risk towards a second predator (Hoverman & Relyea 2008). Because changes in morphological *traits* are not instantaneous, the length of the acclimation period with the first predator could affect the amount of increased predation risk. These temporal changes in interaction modification are the higher-order equivalent of priority effects, and we term them “time-dependent interaction modification.” This term includes all forms of higher-order interactions that can change with species’ arrival times.

In plant communities, such time-dependent interaction modification could arise from plant-soil feedback. Early-arriving plants modify the soil environment, e.g., via allelopathy, changes in the microbiome, and litter addition from previous residents, which reduces the fitness of the later arriving plant (Inderjit & van der Putten 2010; van der Putten *et al*. 2016; Suding & Goldberg 1999). However, the strength of the plant-soil feedback often depends on how long the plant conditions the soil (Ke *et al*. 2021; Lepinay *et al*. 2018). Furthermore, plant-plant interactions can also be mediated by soil microbiome cultivated by a third species arriving before them (Zhang *et al*. 2020). These studies indicate the potential of time-dependent interaction modification in plant communities, but this relationship is still unknown.

To fill this conceptual gap, we explore how time-dependent interaction modifications can arise in nature. We generate population dynamics from a mechanistic plant-soil feedback model that focuses on the interaction between plants and their microbiomes. Then, we apply a framework of time-dependent interaction modification to the simulated data to reveal the relationship between the temporal structure of communities and their higher-order interactions.

## Methods

### Plant-soil Feedback Model

We simulate seasonal population dynamics of plants using a mechanistic, discrete-time model of plant-soil feedback to explore how time-dependent interaction modifications can arise in natural communities (Figure 1). Each year has *T* time steps. We use τ to represent each specific time step within the year (*j* ≤ τ ≤ *T*), and each plant species germinates from dormancy at a specific time step τ = p_x_, where *p*_*x*_ is the germination phenology of the plant x. Note that a “year” defines the suitable growing season for plants and microbiomes, which may be a few months per calendar year. After germination, each plant cultivates microbiomes which affects the mortality rates of all plants (Bever *et al*. 1997). At the end of their life cycles, plants reproduce and senesce. All plants finish their life cycles within a year, regardless of their germination phenology. We assume that life cycles of all plants have identical lengths of *l* time steps (p_x_ + *l* ≤ *T*), such that the plant germinating earliest will also reproduce and senesce earliest in the year.

**Figure 1.**
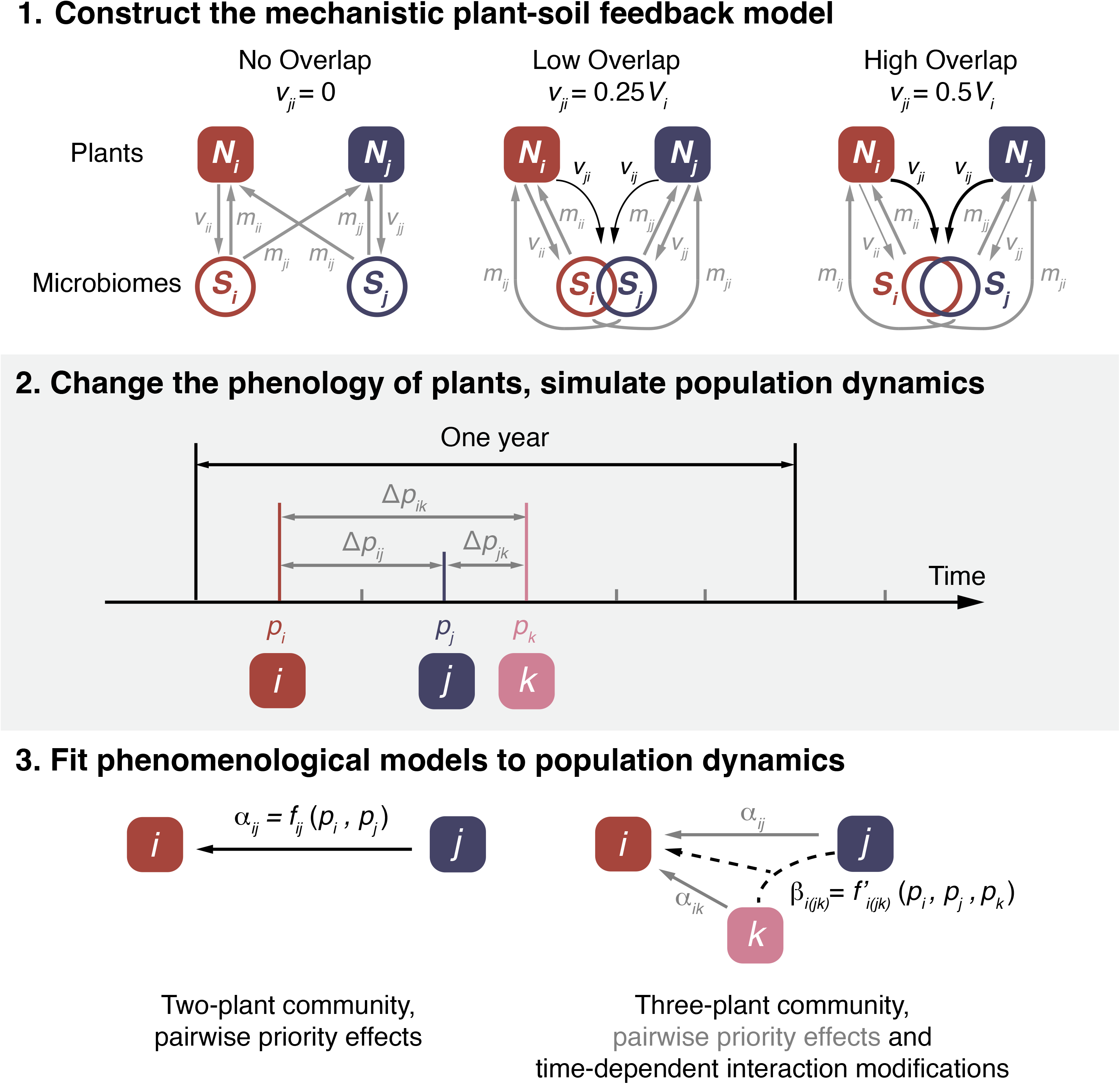
A roadmap of the methods. (1) We first construct a plant-soil feedback model (two-plant model shown for simplicity). With no overlap of associated microbiomes between plants, each plant can cultivate only its associated microbiome (*v*_*ii*_ and *v*_*jj*_), which in turn affects bothplant species (*m*_*ii*_, *m*_*ij*_, *m*_*ji*_, and *m*_*jj*_). With overlap in microbiomes, each plant species can also cultivate the microbiome associated with the other plant species (*v*_*ij*_ and *v*_*ji*_, black arrows); higher overlap leads to higher interspecific cultivation rates. (2) We then apply the above model to communities with two and three plants and their microbiomes to simulate plant population dynamics. Each plant germinates at a certain time, leading to three distinctive relative arrival times, Δp_ij_, Δp_jk_, and Δp_ik_. (3) Finally, we fit phenomenological models to population dynamics generated in (2) to evaluate priority effects and time-dependent interaction modifications. In two-plant communities (*i* and *j*; left), trait-dependent priority effects occur when the per-capita interaction from *j* to *i* (*α*_*ij*_) is determined by the relative arrival time between *j* and *i* (*f*_*ij*_(p_i,_ p_j_). In three-plant communities, the presence and arrival time of plant *k* can determine the interaction between the other two species (β_*i(jk)=*_ *f*_*i(jk)*’_(*p*_*i*,_ *p*_*j*,_*p*_*k*_)), leading to time-dependent interaction modification.

Within the life cycles, the mortality of plant x depends on its density (N_x_(τ)) and the accumulated effects from all microbiomes (Σ_y_ m_xy_ S_y_; Suding *et al*. 2013), where m_xy_ denotes the effect of microbiome associated with plant *y* on plant x, S_*y*_ the density of the microbiome associated with plant y, and d_x_ the density-dependent mortality rate:

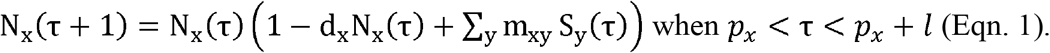

At the end of their life cycle, plants produce seeds (D_x_) with fecundity λ_x_:

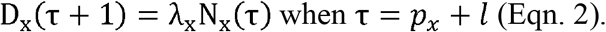

Seeds germinate at the assigned phenology (*p*_*x*_) at the beginning of the next year without additional mortality, i.e., N_*x*_ (τ+ *j*) = *D*_*x*_ (τ) when τ = *p*_*x*_.

To highlight the effect of plant-soil feedback, we focus on interspecific interactions arising from changes in microbiomes (Figure 1) rather than resource competition. This represents a natural scenario where each species is limited by different resources (Tilman 1982).

Within the year, the population of microbiomes (Sx (τ)) increases proportionally with plant density at the rate v_xy_N_y_ once the plants emerge, where v_xy_ is the rate of cultivation by plant y, and is limited by a density-dependent mortality (μ_*x*_S_*x*_ (τ)):

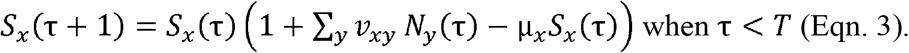

Eqn. 3 specifies that when plants are not present (e.g., before seed germination or after senescence), their associated microbiome immediately starts to decay (Esch & Kobe 2021). We assume a constant proportion (μ_year_) of each microbiome does not carry over to the next year:

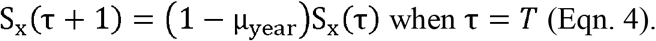

### Time-dependent Interaction Modification

Higher-order interactions are often measured from fitting population data to statistical models (Kleinhesselink *et al*. 2022; Letten & Stouffer 2019). Here, we extend the previous framework of higher-order interactions to define and quantify time-dependent interaction modification in the plant-soil feedback model (Eqn. 1-4). Consider a three-species community that assembles over time (Figure 1), where the change in density (*N*) of a species between time steps can be characterized by a pairwise Ricker model:

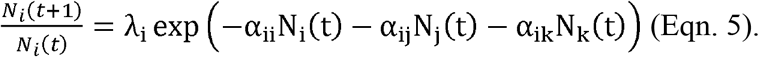

*λ*_*i*_ is the intrinsic growth rate of species *i*, and *α*_*ij*_ is the per-capita effect from species *j* to *i*. In a pairwise system with priority effects (Figure 1), the per-capita interaction from species *j* to *i* can depend on the difference between their germination phenology within a year, *p*_*j*_ − *p*_*i*_, i.e., their phenological difference Δ*p*_*ij*_ (Rudolf 2019; Zou & Rudolf 2023a) (Figure 1). In this case, we can expand Eqn. 5 into:

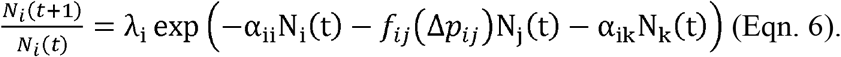

On the other hand, higher-order interactions in this three-species system can be modeled by the following variant of the Ricker model (Kleinhesselink *et al*. 2022):

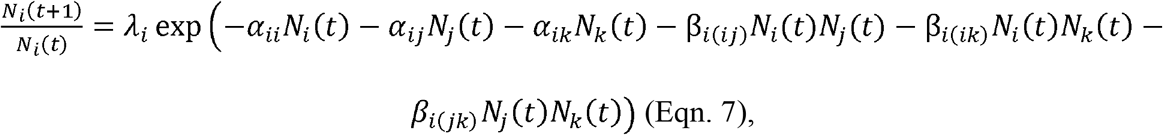

where the β coefficients quantify the strength of the higher-order interaction (HOI), e.g., β_*i(jk)*_ quantifies how densities of species *j* and *k* modify each other’s effect on species *i* (Letten & Stouffer 2019).

If phenological differences between species can affect the strengths of both pairwise and higher-order interactions by changing the corresponding per-capita coefficients (Figure 1), in the most general form, Eqn. 7 can be written as:

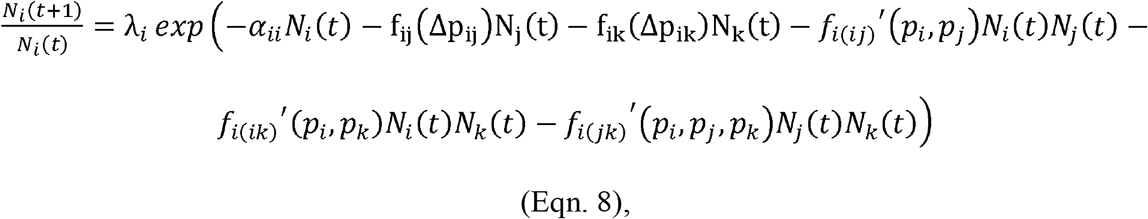

where *f*_*i(jk)*’_′*(p*_*i*,_ *p*_*j*,_ *p*_*k*_) defines how the HOI coefficient, β_*i(jk)*_, depends on the phenology of all three species. If we consider species *i* and *j* as the “focal pair” and fix their phenology while shifting species *k*’s phenology, then *f*_*ij*_ (Δ*p*_*ij*_) and *p*_*j*_ become constants, and β_*i(jk)*_ only depends on *p*_*k*_. We maintain this assumption here to simplify our analyses of three-species communities.

To detect time-dependent interaction modification, we compare *β*_*i(jk)*_ values fitted from population dynamics with different phenology of species *k*(*p*_*k*_). If interaction modifications are absent, *β*_*i(jk)*_ should be close to 0; if *β*_*i(jk)*_ is nonzero but constant for all *p*_*k*_, then the interaction modification does not depend on time. This definition does not consider time-dependent interaction modification via temporal changes in species density, which is beyond our scope.

The accurate quantification of time-dependent interaction modifications requires density gradients of all three species and a temporal gradient of at least one species, a scale rarely feasible for empirical studies. A simplified quantification of time-dependent interaction modifications fits pairwise models between focal species *i* and *j* (e.g., Eqn. 5) to experimental populations with and without species *k*, then examines how the difference of the two pairwise interaction coefficients change with the phenology of species *k* (*p*_*k*_). Under this definition, the time-dependent interaction modification combines the effects of both the density and phenology of species *k*, and its detection requires much less density gradients of all three species (Appendix II). In practice, it simplifies the experimental design but carries stronger assumptions on the community with and without the third species *k*. We discuss the mathematical links between the methods and their pros and cons in Appendix II, but focus here on the first approach (Eqn. 8) to facilitate direct comparison to previous studies (Kleinhesselink *et al*. 2022; Letten & Stouffer 2019). In both approaches, the interpretation of fitted higher-order interactions depends on the specific system and its biological processes, and the statistical power of the phenomenological model structure.

### Simulation

In nature, plants can share pathogenic or mutualistic microbes (Crawford *et al*. 2019; Gilbert & Webb 2007; Sedio & Ostling 2013; Spear & Broders 2021), which can be cultivated by multiple plants in the same community. We simulate this overlap in microbiomes by allowing for the plant x to cultivate not only its associated microbiome but also those associated with other plants (v_yx_ ≠ 0 for all *x* ≠ *y*). Each plant’s total capacity of cultivation (*V*_*x*_ = *v*_*XX*_ + Σ_*y*_ *v*_*yx*_ for all *x* ≠ *y*) is constant, reflecting a finite pool of resources plants can provide to microbiomes. To simulate different levels of shared microbiomes among plants, we partition each plant’s cultivation rate to simulate three scenarios: no overlap (Σ_*y*_ *v*_*yx*_ = 0*V*_*x*_), low overlap (Σ_*y*_ *v*_*yx*_ = 0.25*V*_*x*_), and high overlap in microbiomes (Σ_*y*_ *v*_*yx*_ = 0.5*V*_*x*_). The strength of interspecific cultivation is equally split between microbiomes associated with other plants and therefore depends on the number of plant species. These values ensure that the microbiome always receives the largest (or equal) cultivation effect from its host compared to others.

We focus on the role of soil pathogens only and assume that microbiomes increase the mortality within the year (i.e., m_xy_ < 0; see Table 1), although our methods can be readily extended to mutualistic and mixed microbiomes. We explored the effect of specificity of the pathogens by changing the microbiome’s effect on its host vs. other plants: if *m*_xx_ = *m*_*yx*_, the microbiome is a generalist pathogen to all plants; if |m_xx_| > |*m*_*yx*_| or |m_xx_| < |*m*_*yx*_|, the microbiome is a specialist on its host, or plants other than its host. To simulate the specificity of microbiomes found in nature, we calculated the distribution of *m*_*xx*_/*m*_*yx*_ from an empirical metaanalysis (Yan *et al*. 2022), then randomly drew this proportion to partition the total capacity of feedback. See Appendix I for detailed methods.

**Table 1.** Parameters used for simulations. All values except for phenology are not plant-specific and follow the general notations in Eqn. 1-4. Note that subscripts *x* and *y* can refer to any species, but *i, j*, and *k* refer to specific species.

| Definition | Symbol | Values |
| --- | --- | --- |
| Initial density of plants (as seeds) | $D(0)$ | 1 |
| Initial and baseline density of microbiomes | $S(0)$ | 0.1 |
| Length of the year | $T$ | 12 |
| Length of plants' life cycle | $l$ | 6 |
| Plant mortality rate within the life cycle | $d$ | 0.1 |
| Microbiome mortality rate within the year | $\mu$ | 0.1 |
| Microbiome mortality at the end of the year | $\mu_{year}$ | 0.7 |
| Plant fecundity | $\lambda$ | 3 |
| Total capacity of cultivation (plants to microbiomes) | $V$ | 0.3; or drawn from uniform distribution (0.1, 0.3) |
| Cultivation rate from plant $x$ to its associated microbiome | $v_{xx}$ | $V_x$ if no overlap, $0.75V_x$ if small overlap, $0.5V_x$ if high overlap |
| Cultivation rate from plant $x$ to other plants' associated microbiome | $v_{yx}$ for all plants $x \neq y$ | 0 if no overlap, $0.25V_x/n$ if small overlap, $0.5V_x/n$ if high overlap; $n$ is the number of plant species other than itself |
| Effects of microbiomes on plants | $m_{xy}$ for all plants $x$ and $y$ | $-0.1$ or $-0.3$; or drawn from uniform distribution ($-0.5$, $-0.1$) |
| Phenology of plants $i$, $j$ and $k$ | $p_i, p_j, p_k$ | 1 (earliest) to 6 (latest) |
| Phenological difference between the focal pair $i$ and $j$ | $\Delta p_{ij}$ | $-5$ to $5$ |

Each year has 12 time steps (*T*= 12) and the length of plant life cycles (*l*) is 6 time steps (*l*= 6); therefore, the earliest germination phenology is 1 (the beginning of the year), and the latest 6. Plants’ germination phenology does not change over years or with other plants. This range ensures enough time for species interaction while allowing for a proper gradient of phenological differences between plants. We choose identical fecundity λ and effects of microbiomes *m*_*xy*_ (Table 1) for all plants, such that any differences in population dynamics would arise from phenological differences between plant species, different overlaps of microbiomes, and different host specificity of microbiomes.

To accurately obtain interaction coefficients (Eqn. 8), we used a response surface design (commonly used in empirical studies; Inouye 2001; Kleinhesselink *et al*. 2022) with 6 densities of each plant species, yielding 36 and 216 combinations of initial densities for two-plant and three-plant communities. We ran all response surface simulations for one year, representing the typical duration of plant experiments. We simulate two-plant communities with a gradient of 11 phenological differences from Δ*p*_*ij*_ = *p*_*j*_ − *p*_*i*_ = −5 (species *j* early by 5 time steps) to Δ*p*_*ij*_ = 5 (species early by 5 time steps). In three-plant communities, when a third plant is added to the focal pair, it could germinate either before (scenario I), between (scenario II), or after the focal species pair (scenario III; Figure 3). For simplicity, we examined scenarios I and III by fixing the phenology of the focal pair (*i* and *j*) both at the latest (time 6; scenario I) or at the earliest (time 1; scenario III) and changed the phenology of the third plant *p*_*k*_ ranging from 1 to 6. This allows all plants to germinate simultaneously in some cases. To explore the case where plant *k* germinates between the focal pair (scenario II), we let plant *i* germinate at the earliest (time 1) and *j* at the latest (time 6) and again let 1 ≤ *p*_*k*_ ≤ 6.

We fit a discrete-time Ricker model containing all higher-order interaction terms to plant population at the end of the year (Eqn. 8). The Ricker model structure fits our simulated data best among several other discrete-time population models (Hart *et al*. 2018) based on Akaike Information Criterion (AIC; Appendix I, Figure S1). To evaluate whether a model with higher-order coefficients indeed significantly improves model performance, we fitted three-plant population dynamics to pairwise Ricker models (Eqn. 1) with either plants *i* or *j* as the focal species, then compared both the root-mean-squared error (RMSE) and the AIC values of the pairwise (Eqn. 1) and higher-order Ricker model (Eqn. 8) at different *p*_*k*_. Finally, to test the sensitivity to model parameters, we randomly drew 100 sets of microbial effects (*m*) and cultivation rate (V) from a uniform distribution (Table 1), and fitted Eqn. 8 to obtain a distribution of higher-order coefficients.

We run the model for 50 years (600 time steps) with equal initial densities of plants and microbiomes to evaluate coexistence; the duration ensures that all populations reach equilibrium.

We conducted all simulations in R 4.2.1 (R Core Team 2022) and fitted models using the function *nlsLM()* in the R package *minpack*.*lm* (Elzhov *et al*. 2016). The code is available at https://doi.org/10.5061/dryad.b8gtht7kf.

## Results

### Two-plant Communities

We first examine two-plant communities to analyze the causes and consequences of priority effects. The late plant always has a lower population, and this disadvantage is smaller with smaller differences between the germination phenology of the two plants (Figure 2A). The effect of the early plant on the late plant increases when it germinates earlier. Changes of *α*^*ij*^ and *α*^*ji*^ with differences in germination phenology Δ*p*_*ij*_ for both plants are symmetrical (Figure 2B) and nonlinear. Note that when one plant germinates very late (|Δ*p*_*ij*_| ≥4), its effect on the early plant is negligible because the temporal overlap between the two plants is too short, and the effect it experiences from the early plant slightly decreases because the early plant’s microbiome declines without its host.

**Figure 2.**
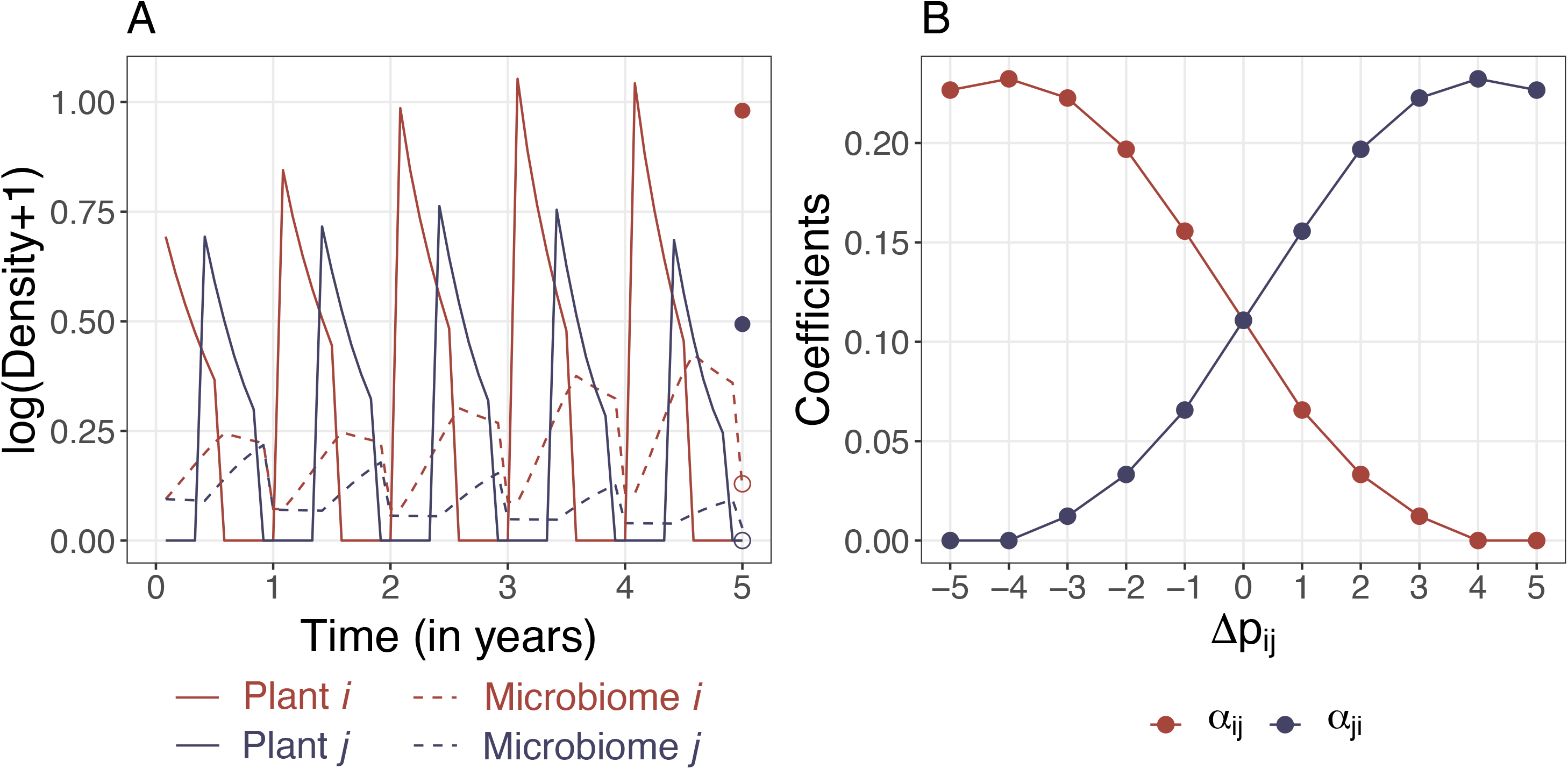
Results of two-plant communities with equal microbial effects (|m_xx_| = |*m*_*yx*_|). A. Population dynamics of the plant-soil feedback model with two plants showing a clear early-arriver advantage (Δ*p*_*ij*_ = 4). Densities of plants and microbiomes are in solid and dashed lines. Equilibrium densities (after 50 years) of plants and microbiomes are shown in solid and empty dots. Dynamics of seeds are not shown. B. Fitted interspecific interaction coefficients between plants *i* and *j* along a temporal gradient of relative arrival times Δ*p*_*ij*_, . A Δ*p*_*ij*_ < 0 means that *j* arrives early, Δ*p*_*ij*_ > 0 means that arrives early.

The overlap between plants’ microbiomes interacts with the specificity of microbiomes to affect the final populations and interaction coefficients. When the microbiome affects other plants more negatively than its host (|m_xx_| < |*m*_*yx*_|), for *x* ≠ y), each plant receives less feedback from its microbiome, and the early plant tends to exclude the late plant. However, with increasing overlap the early plant cultivates the late plant’s microbiome, which subsequently exerts negative feedback to the early plant, promoting coexistence (Figure S5). When the microbiome affects its host the most (|m_xx_| > |*m*_*yx*_|), increasing overlap leads to a stronger negative interaction between plants and promotes monodominance (Figure S6).

### Three-plant Communities: Population Dynamics

We examine patterns of equilibrium population density following three scenarios of plant phenology (plant *k* germinates before, between, or after the focal pair; Figure 3). In general, the earlier a plant germinates relative to its competitors, the higher its equilibrium population (Figure 3). However, when all three plants germinate at different times (scenario II), both the earliest and the last plants benefit from the later germination of the intermediate species, while only the intermediated species itself is negatively affected. This alternating pattern indicates a time-dependent interaction chain: When plant *k* germinates later than plant (scenario II), plant experiences lower total microbiome densities and reaches a higher equilibrium population. Because plant *k* germinates later it has also less time to cultivate its microbiome. This combination of reduced density and cultivation time of the intermediate plant (*k*) results in a decline in the total density of the microbiome, which benefits the last plant *j* (Figure S3).

**Figure 3.**
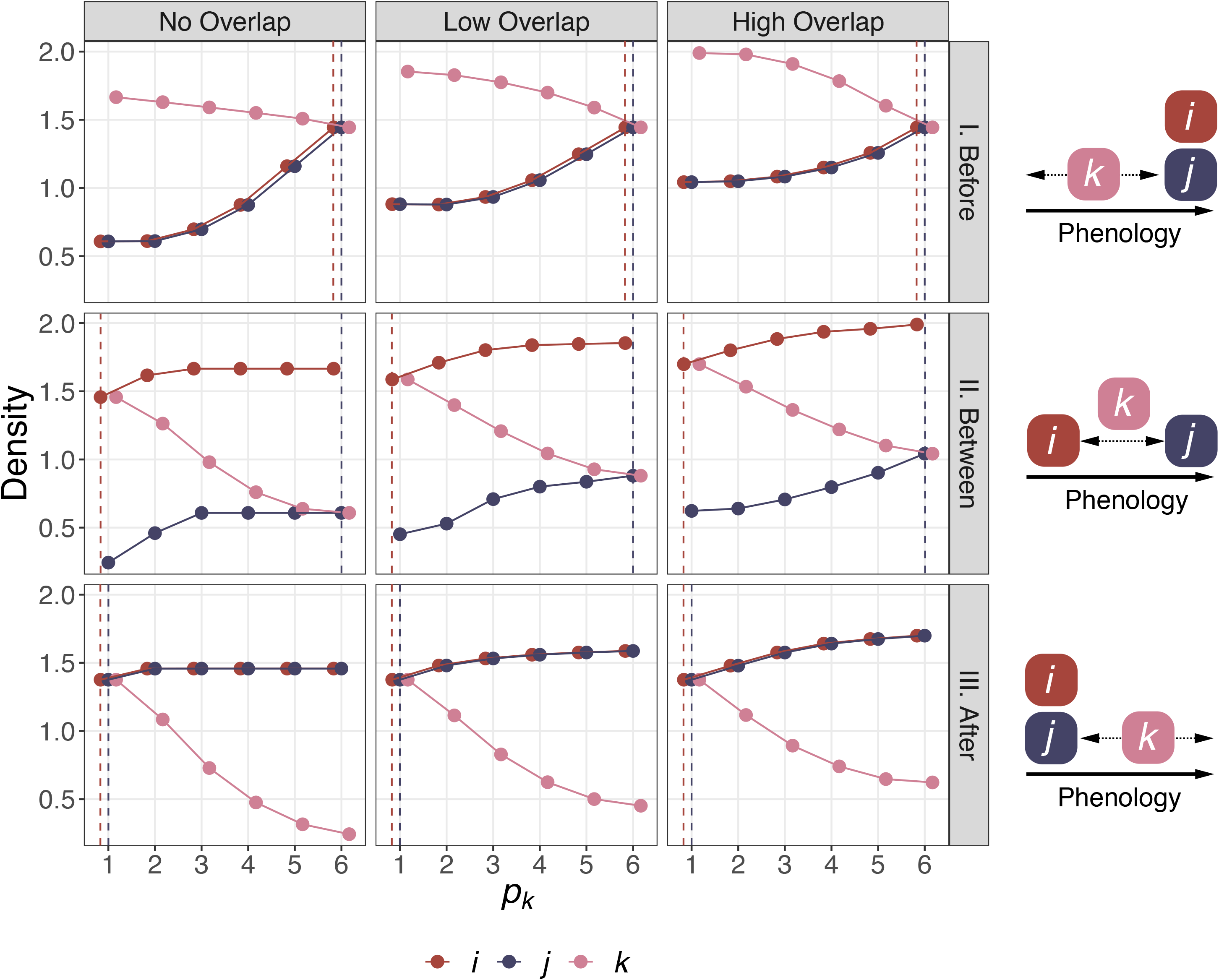
Equilibrium population (after 50 years) of three plant species in three-plant communities with equal microbial effects (|m_xx_| = |*m*_*yx*_|). Vertical red and blue dashed lines indicate the germination phenology of plants and *j*. Scenario I shows plant *k* germinating before the focal pair *i* and *j*, Scenario II shows *k* germinating between the focal pair, and Scenario III shows *k* germinating after the focal pair. When species arrive at the same time, their equilibrium population is identical; points and lines of each plant are jittered for visibility.

Like in two-plant communities, levels of overlap in microbiome cultivation among plants interact with the specificity of microbiomes. When all microbial effects are equal, larger overlaps (higher inter-vs. intraspecific cultivation) lead to higher equilibrium populations because plants with earlier phenology devote less to cultivating their microbiomes, leading to a slower increase of microbiomes before the late plants germinate and subsequently a lower overall density of microbiomes (Figure S3). Larger overlap promotes coexistence if the microbial effect on its host is smaller than to others (|m_xx_| < |*m*_*yx*_|)but promotes monodominance if (|m_xx_|> |*m*_*yx*_|) (Figure S7-S8).

### Time-dependent Interaction Modifications

We examine patterns of time-dependent interaction modifications using the higher-order coefficient associated with densities of all three plants *β*_*i(jk)*_ for plant *i β*_*i(jk)*_ for plant *j*, and *β*_*i(jk)*_ for plant *k*) following the same scenarios of plant phenology (Figure 4). Models that include higher-order interaction terms almost always fit best, except when fitted higher-order coefficients are close to 0, despite additional parameters (Figure S16-S17). These results support the observed time-dependent interaction modifications quantified by our methods. The fitted higher-order coefficients are generally smaller in value than pairwise coefficients, but the comparison of the strength of higher-order versus pairwise interaction is contingent on species densities (Eqn. 7).

**Figure 4.**
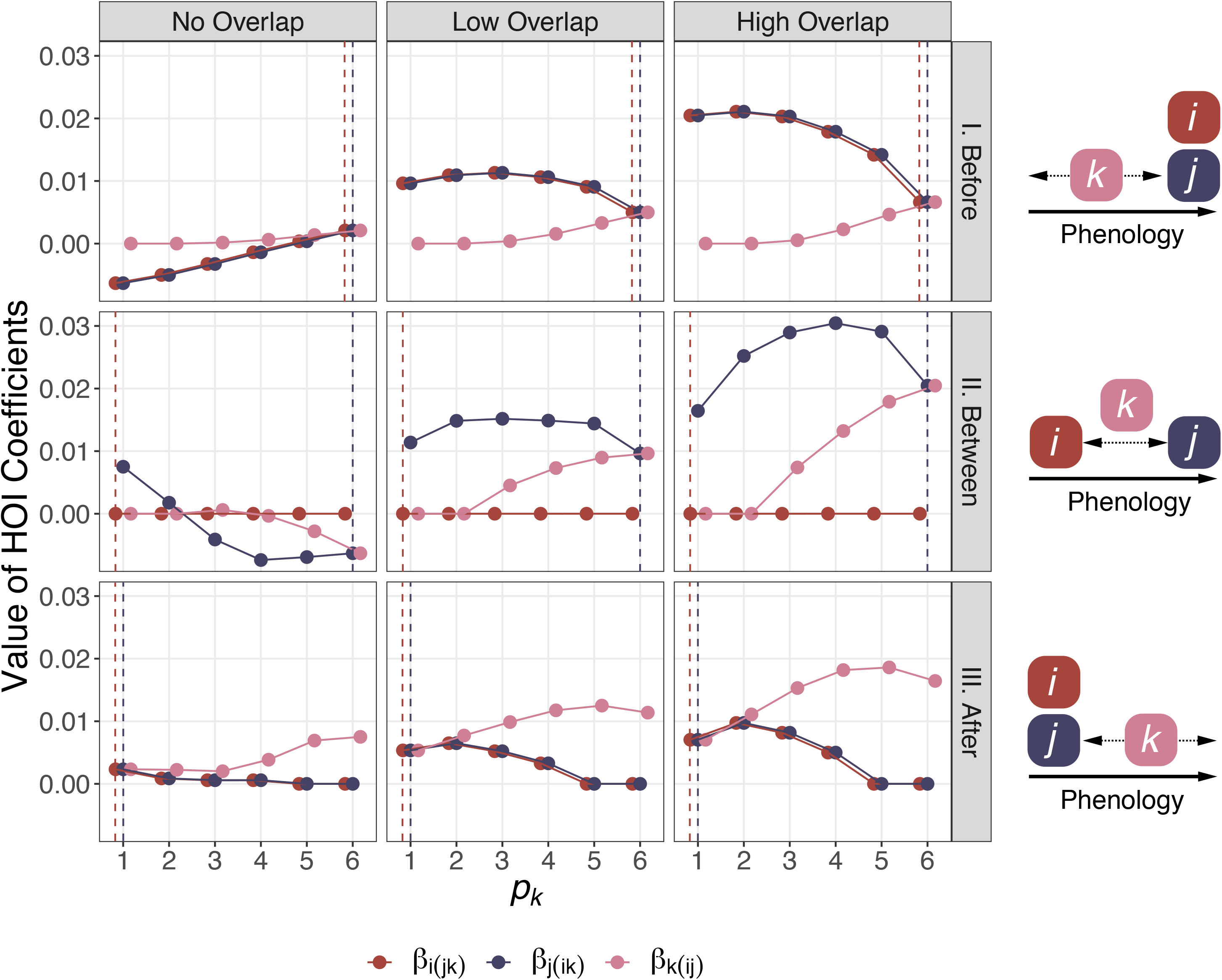
Time-dependent interaction modification in three-plant communities with equal microbial effects (|m_xx_| = |*m*_*yx*_|). The higher-order interaction coefficients *β*_i(jk)_ and *β*_i(jk)_ are fitted according to Eqn. 8. Vertical red and blue dashed lines indicate the germination phenology of plants *i* and *j*. Scenario I shows plant *k* germinating before the focal pair and *j*, Scenario II shows *k* germinating between the focal pair, and Scenario III shows *k* germinating after the focal pair. When the focal pair germinates at the same time (Scenarios I and III), the two plants are equivalent, and higher-order interaction coefficients *β*_i(jk)_ and *β*_i(jk)_ are identical; points and lines of each coefficient are jittered for visibility.

In general, all higher-order interaction coefficients depend on the relative phenology of plants and the level of overlap in associated microbiomes (Figure 4). When plant *k* germinates before both plants *i* and *j* (the focal pair; scenario I, the strengths of higher-order interactions experienced by late plants and *j* (*β*_*i(jk)*_ and *β*_*i(jk))*_ decreases with earlier germination phenology of plant *k* without overlap, increases then decreases with low overlap, and decreases nonlinearly with high overlap of microbiomes. Without overlap in microbiomes, plants *i* and *j*’s microbiomes are not cultivated by early-germinating plant *k*, and the focal pair’s exposure to plant *k*’s microbiome also decreases as plant *k* germinates earlier. The combined effects lead to negative *β*_*i(jk)*_ and *β*_*i(jk)*_ values, weakening the *overall* interactions experienced by plants *I* and *j*.

When plant *k* germinates between plants *i* and *j* (scenario II), the first plant always experiences the weakest higher-order interaction. Similar to scenario I, without overlap in microbiomes, the latest-germinating plant *j* is less affected by other plants’ microbiomes, while the cascading effect between the three plants observed in population dynamics further weakens the interaction it experiences. By germinating later, plant *k* decreases its temporal overlap with the earliest plant and its microbiome. These effects are reflected by the negative *β*_*i(jk)*_ and *β*_*i(jk)*_ values. With higher overlap in microbiomes, the early-germinating plants cultivate microbiomes of late-germinating plants, which experience stronger higher-order interactions.

When plant *k* germinates after both plants and *j* (scenario III), higher-order interactions experienced by plants *i* and *j* (*β*_*i(jk)*_ and *β*_*i(jk)*)_ generally decrease with later phenology of plant *k* because its effect on the microbiomes associated with the focal pair diminishes, while the higher-order effect of the early plants on the late germinating plant *k* (*β*_*i(jk)*)_ generally increases because the effects from the focal pair get stronger when they germinate relatively earlier.

Although microbiomes’ specificity (i.e., the strength of m_xx_ vs. m_yx_) interacts with the overlap of microbiomes to determine the long-term coexistence between three plants, qualitative patterns of time-dependent interaction modifications are unchanged (Figure S9-S10). Drawing microbial effects and cultivation rates from uniform distributions, or from the distribution informed by the range found in experiments (where *m*_*xx*_/*m*_*yx*_ ∼*lognormal* (0.109,1.168); see Appendix I) changes the strengths but not patterns of time-dependent interaction modifications (Figure S12-S15).

We focus on terms that involve all three plants (*β*_*i(jk)*_, *β*_*j(ik)*_ and *β*_*k(ij)*_) because they are unique to three-plant communities. All other higher-order terms in Eqn. 8 change with the phenology of plant *k*, and both the magnitudes and the temporal patterns shift along a continuum with increasing overlap in microbiomes (Figure S11).

## Discussion

To understand the present and future of ecological communities, we need to first understand their history (Fukami 2015; Gause 1934). Here, we bring this temporal perspective to multispecies communities with higher-order interactions. Using a mechanistic model of plant-soil feedback, we show that the germination phenology of a third plant determines how it modifies interaction between other plants. The strength of this interaction modification depends on the overlap in associated microbiomes between all plants which control indirect interaction effects in this system. Together, these results highlight the time-dependence of higher-order interactions in complex communities and provide a road map to further exploring the temporal dimension of species interactions in diverse communities.

Our framework puts higher-order interactions in a temporal context. In our model, the accumulation of microbiomes over time affects the strength of plant-plant interactions. The germination phenology of a third plant marks the starting point of its microbiome cultivation, subsequently the total microbiome densities of the community and their effects on plants. Importantly, the interaction modification changes with phenological differences because the underlying biological process is not instantaneous, despite models of community dynamics that often assume otherwise. Non-instantaneous processes in real ecosystems can lead to time lags between different populations, especially when their life cycles are of similar time scales (e.g. the hare-lynx cycle; Nisbet 1997; Stenseth *et al*. 1998; Adler *et al*. 2008). The ubiquity of these temporal processes implies that all higher-order interactions have the potential to be time-dependent.

Higher-order interactions can arise from various ecological mechanisms. When one species alters the environment, the modification can often affect how other species interact, leading to higher-order interactions (Wootton 1993; Zhang *et al*. 2020). When these environmental modifications progress with time, they can also lead to priority effects (Chappell *et al*. 2022). The temporal process of environmental modification is therefore a potential source of time-dependent interaction modification in complex communities. In our model, the cultivation of microbiomes by plants modifies the fitness of other plants, and the different germination phenology affects the temporal extent of these environmental modifications, leading to time-dependent interaction modifications. Similarly, one species can alter the traits of other species, which can also progress with time, leading to time-dependent interaction modifications (Hoverman & Relyea 2008; O’Keeffe *et al*. 2021; Werner & Peacor 2003). Theoretical studies often mechanistically model individual species’ demographic parameters or traits that give rise to higher-order interactions, such as growth rates (Kleinhesselink *et al*. 2022; Lai *et al*. 2022), fecundity (Mayfield & Stouffer 2017), and size (Levine *et al*. 2017). Since these traits often change with species’ arrival times (Blackford *et al*. 2020; Rasmussen *et al*. 2014) the strengths of resulting higher-order interactions should also be time-dependent. Incorporating time-dependence in these traits is essential to understand changes in higher-order interactions over time in complex communities.

In our model, the interaction between two generalist-specialist gradients not only determines the magnitude of higher-order interactions but also the coexistence of plants. A plant can specialize in cultivating its microbiome (no overlap), or cultivate all other plants’ microbiomes (total overlap), whereas a microbiome (in our model, pathogenic) can specialize in its host or other plants (higher microbial effects), or equally affect all plants (uniform microbial effects). If a microbiome specializes in plants other than its host, its host would effectively have lower self-limitation than limitation to other plants, leading to monodominance. However, higher overlap allows it to limit itself by cultivating other plants’ microbiomes promoting coexistence. In this case, higher overlap promotes both time-dependent interaction modifications and coexistence. On the other hand, when a microbiome specializes on its host, higher overlap promotes monodominance by increasing the microbiomes of other plants, raising the limitation to others above that of the host plant. These dynamics arising from the two pairs of generalist vs. specialist gradients provide a mechanistic explanation of the strengths of higher-order interactions and coexistence between plants, but not necessarily a direct relationship between the two. Our results provide a detailed perspective on how the composition of generalist vs. specialist can shape dynamics across communities (Garzon-Lopez *et al*. 2015; Holt & Lawton 1993; Jiao & Cortez 2022; Snyder & Ives 2001). Although the temporal aspect of these indirect interactions is studied in the context of pairwise priority effects (Clay *et al*. 2019; Jiao & Cortez 2022), future studies need to consider the role of time-dependent interaction modifications in these intrinsically complex communities (e.g., O’Keeffe *et al*. 2021).

In nature, plants often share mutualists or pathogens, especially among those more phylogenetically related (Davison *et al*. 2015; Gilbert & Webb 2007; Põlme *et al*. 2018), and the sharing is often assumed in studies that explicitly model mutualists and pathogens (Abbott *et al*. 2021; Jiang *et al*. 2020). Although the net antagonistic effect of soil microbes can be common (and assumed in our model; Lekberg *et al*. 2018), mutualistic effects from microbiomes can change niche or fitness differences between plants (Kandlikar *et al*. 2019; Ke & Wan 2020; Yan *et al*. 2022), affecting population dynamics and fitted interaction coefficients. The pathogenic vs. mutualistic roles of microbiomes can further interact with the generalist-specialist gradients in our model (Semchenko *et al*. 2022). For instance, specialist mutualists can amplify fitness differences between plants and promote monodominance or positive frequency dependence (Xi *et al*. 2021); more abundant plants can be more exposed to their specialist pathogens, leading to conspecific negative density dependence that favors biodiversity (Hülsmann *et al*. 2021; Liang *et al*. 2016; Sedio & Ostling 2013). Studying the specific roles of microbiomes will greatly improve our understanding of pairwise and higher-order interactions between plants and mechanisms of coexistence in highly diverse communities.

Increasing evidence suggests that the plant-soil feedback is a highly dynamic process, challenging the classic assumption that microbial dynamics are much faster than plant dynamics such that feedback between plant and microbiome is instantaneous (the “separation of time scales”; Eppinga *et al*. 2018; Kandlikar *et al*. 2019; Mack *et al*. 2019; Ke & Wan 2020). We model the change of microbial community compositions over the same time scale as plants and the decay of plant-associated microbiomes after the senescence of the host. In nature, these common processes can lead to rich dynamics of microbiomes and strongly affect the strengths of plant-soil feedback (Chung 2023; Dombrowski *et al*. 2017; Esch & Kobe 2021; Hannula *et al*. 2019, 2021; Ke *et al*. 2021; Ke & Levine 2021; Lepinay *et al*. 2018; Tan *et al*. 2021), and in our model, they lead to temporal changes in pairwise and higher-order interactions between plants.

Life histories of plants and soil microbes vary greatly across ecosystems. Several biologically realistic assumptions in our model may therefore only apply to certain communities. We model large differences in germination phenology, which represents systems where native and invasive species respond differently to environmental cues (Cleland *et al*. 2015; Wainwright & Cleland 2013). In some annual plant communities where germination is highly synchronized (e.g., California annual grassland; Chiariello 1989; Bart *et al*. 2017), the potential for priority effects and time-dependent interaction modifications may be reduced. We assume fixed lengths of life cycles among plants. However, the plasticity of life history could lead to novel tradeoffs between species, affecting the presence and strengths of higher-order interactions (Kleinhesselink *et al*. 2022; Levine *et al*. 2022). We also assume no mortality or dormancy of seeds. Including a seed bank can promote coexistence by buffering plants against competition or environmental stress (Lennon *et al*. 2021), subsequently affecting the strength of time-dependent interaction modifications. Finally, we assume that all microbiomes are present at the same density before plants germinate. This fixed baseline does not consider the potential differences in the life cycles and phenology of soil microbiomes (Rudgers *et al*. 2020) or the potential for early-germinating plants to associate preferentially with early microbes. Nevertheless, our model provides an important baseline for future empirical and theoretical works on the temporal dynamics of complex plant communities.

Our framework of time-dependent interaction modification lays the groundwork for quantifying temporal changes pairwise and higher-order interactions in multispecies communities, advancing the understanding of their assembly over time (Song *et al*. 2020, 2021b, a). However, our results do not directly address the link between higher-order interactions and coexistence (e.g., Bairey *et al*. 2016; Buche *et al*. 2023; Gibbs *et al*. 2022; Grilli *et al*. 2016).

Resolving higher-order interactions along the temporal axis reveals new processes that could stabilize or destabilize communities, such as fluctuations of arrival times (phenological variations) over years (Carter *et al*. 2018; Rudolf 2019; Theobald *et al*. 2017; Zou & Rudolf 2023b), further contributing to the context-dependency of the complexity-stability relationship. The concept of time-dependent interaction modification combines the two burgeoning fields, higher-order interactions, and time-explicit ecology. This temporal layer of realism is essential to understanding the processes shaping complex communities in nature, and to predicting how they will change as climate change continues to reshuffle species phenology worldwide.

## Author Contributions

H.-X.Z. conceived the idea, developed the methods, performed the simulation, and analyzed the results with assistance from X.Y. and V.H.W.R. H.-X.Z. and X.Y. designed the plant-soil feedback model. H.-X.Z. wrote the first draft and all authors contributed to revisions.

## Acknowledgments

We thank Po-Ju Ke, Gaurav Kandlikar, and Tom E.X. Miller for comments on the manuscript and members of the Levine Lab at Princeton University for thoughtful discussions. Comments from the editor and three anonymous reviewers greatly improved the manuscript. Funding was provided by NSF DEB-1655626. X.Y. is supported by the Stengl-Wyer Fellowship.

