## Supplementary Material for "Time-dependent Interaction Modification Generated from Plant-soil Feedback"

### Appendix I: Supplementary Results

#### *Model Selection*

The statistical detection of higher-order interaction strongly depends on the base model structure that is fit to the data (Letten and Stouffer 2019, Kleinhesselink et al. 2022). We test four commonly used discrete-time phenological models that contain four parameters of population growth ($\lambda$), intraspecific ($\alpha_{ii}$), and interspecific interaction coefficients ($\alpha_{ij}$): Lotka-Volterra, Beverton-Holt, Ricker, and Law-Watkinson (Eqn. S1-S4); Hart et al. 2018:

$$\frac{N_i(t+1)}{N_i(t)} = \lambda_i(1 - \alpha_{ii}N_i(t) - \alpha_{ij}N_j(t)); \tag{S.1}$$

$$\frac{N_i(t+1)}{N_i(t)} = \frac{\lambda_i}{1 + \alpha_{ii}N_i(t) + \alpha_{ij}N_j(t)}; \tag{S.2}$$

$$\frac{N_i(t+1)}{N_i(t)} = \lambda_i \exp(-\alpha_{ii}N_i(t) - \alpha_{ij}N_j(t)); \tag{S.3}$$

$$\frac{N_i(t+1)}{N_i(t)} = \frac{\lambda_i}{1 + N_i(t)^{\alpha_{ii}} + N_j(t)^{\alpha_{ij}}}. \tag{S.4}$$

We generate example data of a two-plant community with low overlap in microbiome ($v_{yx} = 0.2V_x$; see Table 1) simulated at different germination phenology of the two plants ($-5 \leq \Delta p_{ij} \leq 5$) using the standard fecundity, mortality, microbial effects, and cultivation rates as in Table 1. We fit the above four phenological models to the data, then calculate the Akaike Information Criterion (AIC) of each fitted model. Because the two plant species are equivalent except for their different phenology, here we show results where plant $i$ is the focal species. The Ricker model (Eqn. S3) has the lowest AIC value except when plant $i$ germinates very early ($\Delta p_{ij} \geq 3$), where the Law-Watkinson model (Eqn. S4) has the lowest AIC value (Figure S1). We therefore fit Ricker model to all data generated from two- and three-plant communities to ensure all fitted coefficients are statistically comparable.

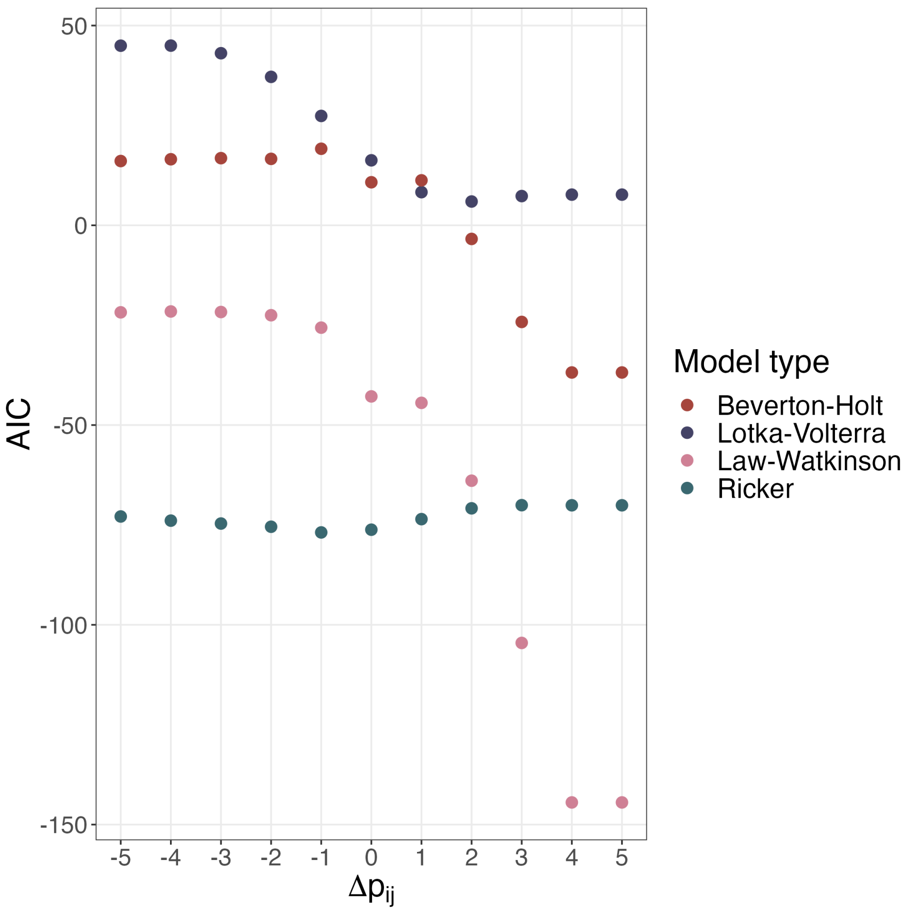

Figure S1: AIC values of four phenological models fitted to simulated plant data. The lower the value, the better the model fit. A $\Delta p_{ij} < 0$ means that j arrives early, $\Delta p_{ij} > 0$ means that i arrives early.

#### *More Details on Two-plant Communities*

Here, we elaborate on details of two-plant communities using equal effects from the microbiomes ($m_{xx} = m_{yx}$). With no overlap in the microbiomes across plants, the density of the microbiome associated with the early plant grows faster and reaches a higher maximum within a year because the plant does not divert its resources to other microbiomes, but it also declines faster because after its host senesces, it cannot be cultivated by the later plant (Figure 2B). With overlap in microbiomes, the fixed total capacity of cultivation reduces growth rates of the plant's associated microbiome, leading to a smaller peak density of the early plant's microbiome, but a higher density of the late plant's microbiome (Figure S2A). Together, these temporal dynamics lead to approximately the same densities of microbiomes with different levels of overlap. When the overlap in the microbiome is less than 50% (i.e., $0 \leq v_{ji} < v_{ii}$), the microbiome associated with the early plant reaches higher density; when the intraspecific cultivation rate is equal to the interspecific cultivation rate ($v_{ji} = v_{ii}$), population dynamics of the two microbiomes are identical. Interactions between plants are generally weaker without overlap in microbiomes because the late plant experiences weaker effects from the early plant's microbiome compared to scenarios with higher overlap in the microbiomes within the first year (Figure S2A). Interaction-phenology functions with small and large overlaps are the same because population dynamics of plants are identical in this scenario but can be quantitatively different with alternative $v_{ii}$ and $v_{ji}$ values (see, for example, Figure S5-S6).

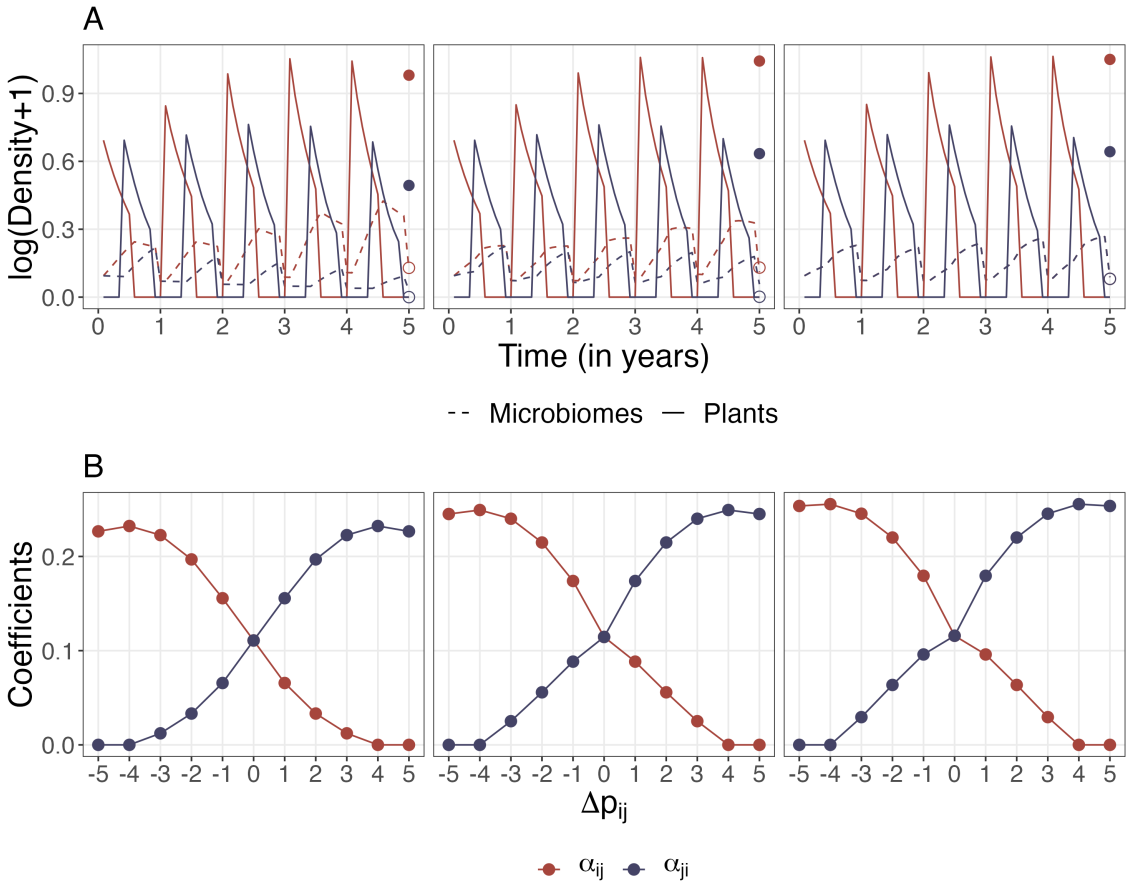

Figure S2: Two-plant communities with identical effects from microbiomes to plants. From left to right: no, medium, and high overlap in associated microbiomes. A. Population dynamics of two-plant communities. Equilibrium densities (after 50 years) of plants (in seeds) and microbiomes at the end of the year are shown in solid and empty dots. B. Fitted interspecific interaction coefficients.

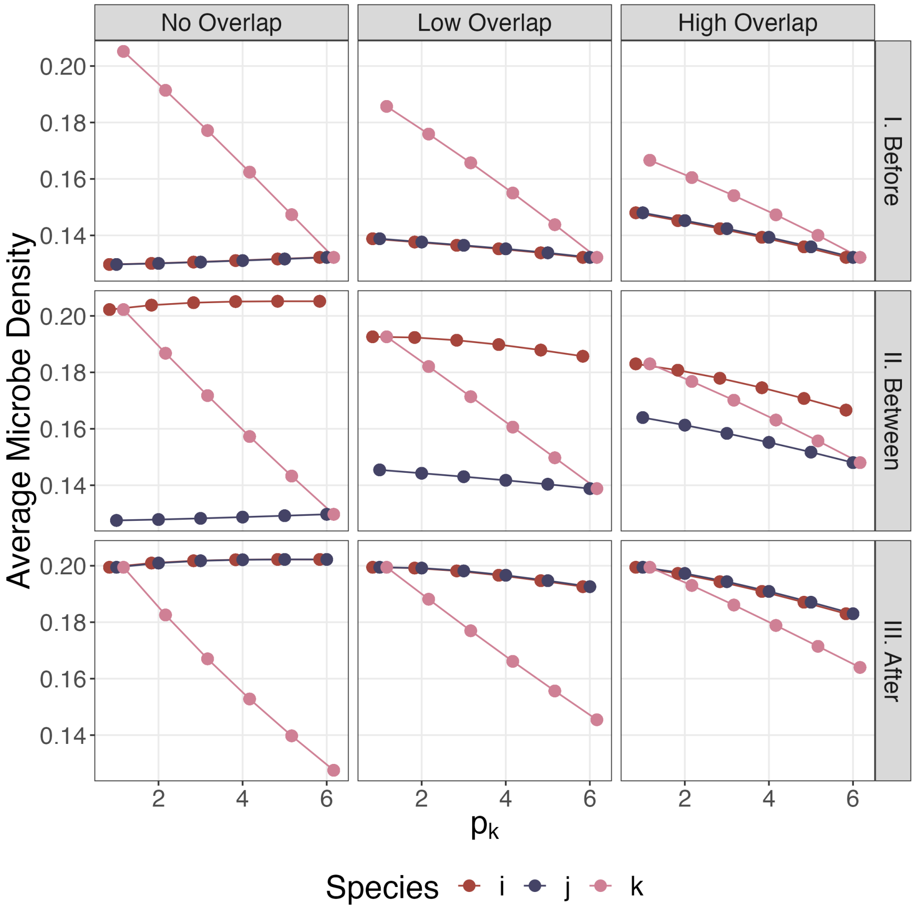

Figure S3: Average densities of microbiomes per time step of the first year change with the phenology of plant $k$. We show the dynamics in the first year because we only fit the plant densities after the first year to derive phenomenological interaction coefficients.

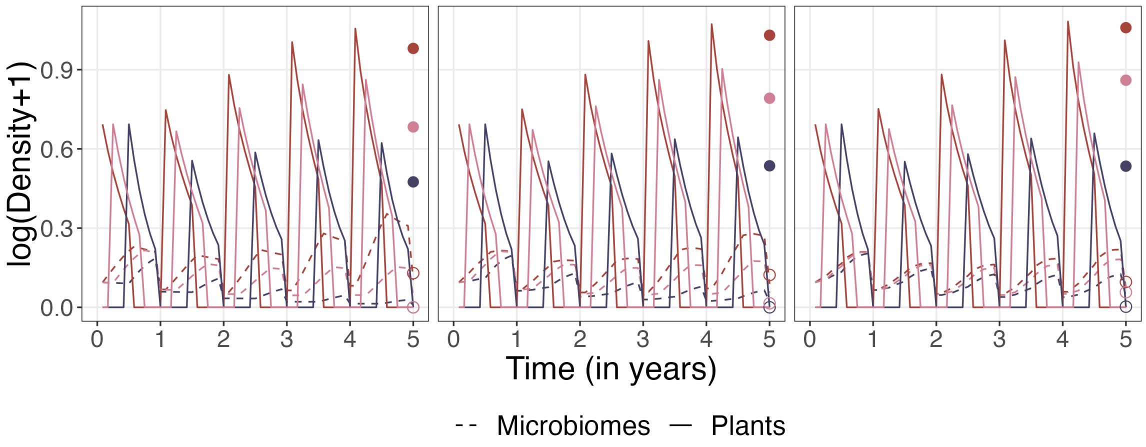

Figure S4: Population dynamics of three-plant communities showing a clear early-arriver advantage. Densities of plants and microbiomes are in solid and dashed lines. Equilibrium densities (after 50 years) of plants and microbiomes are shown in solid and empty dots.

#### *Microbiome Specificity and Coexistence*

We change the relative strengths of the microbial effect to its host, $m_{xx}$, and other plants $m_{yx}$ (for all $y \neq x$). The standard scenario presented in the main text uses $m_{xy} = -0.3$ for all $x$ and $y$. To explore the case where the microbiome is a specialist pathogen of its host, we use $m_{xx} = -0.3$, $m_{xy} = -0.1$; to explore the case where the microbiome preferably lower other plants' survival, we use $m_{xx} = -0.1$, $m_{yx} = -0.3$.

##### *Two-plant Community*

The specificity of microbiomes strongly affects fitted interaction coefficients and species coexistence. When $|m_{xx}| < |m_{yx}|$, the early plant excludes the late plant at no or low overlap in microbiomes, but the two plant coexist with higher overlap (Figure S5). The fitted interaction coefficients also decrease with higher overlap. This is because with $|m_{xx}| < |m_{yx}|$, the early-arriving plant $x$ limits others more than itself by the stronger negative feedback of its microbiome to other plants. But with higher overlap, plant $x$ can also cultivate other plants' microbiomes, which can in turn limit the growth of plant $x$ itself.

On the other hand, when $|m_{xx}| > |m_{yx}|$, higher overlap increases interaction strengths between plants and promote monodominance (Figure S6). As overlap increases, the early-arriving plant $x$ cultivates more microbiome of the late plant, which exerts negative feedback to the late plant. Although we do not observe monodominance with the range of parameters examined (monodominance is difficult because in this case, each plant limit itself more than the other), the difference in the two plants' equilibrium population increases with higher overlap, meaning that the late plant is increasingly likely to be excluded.

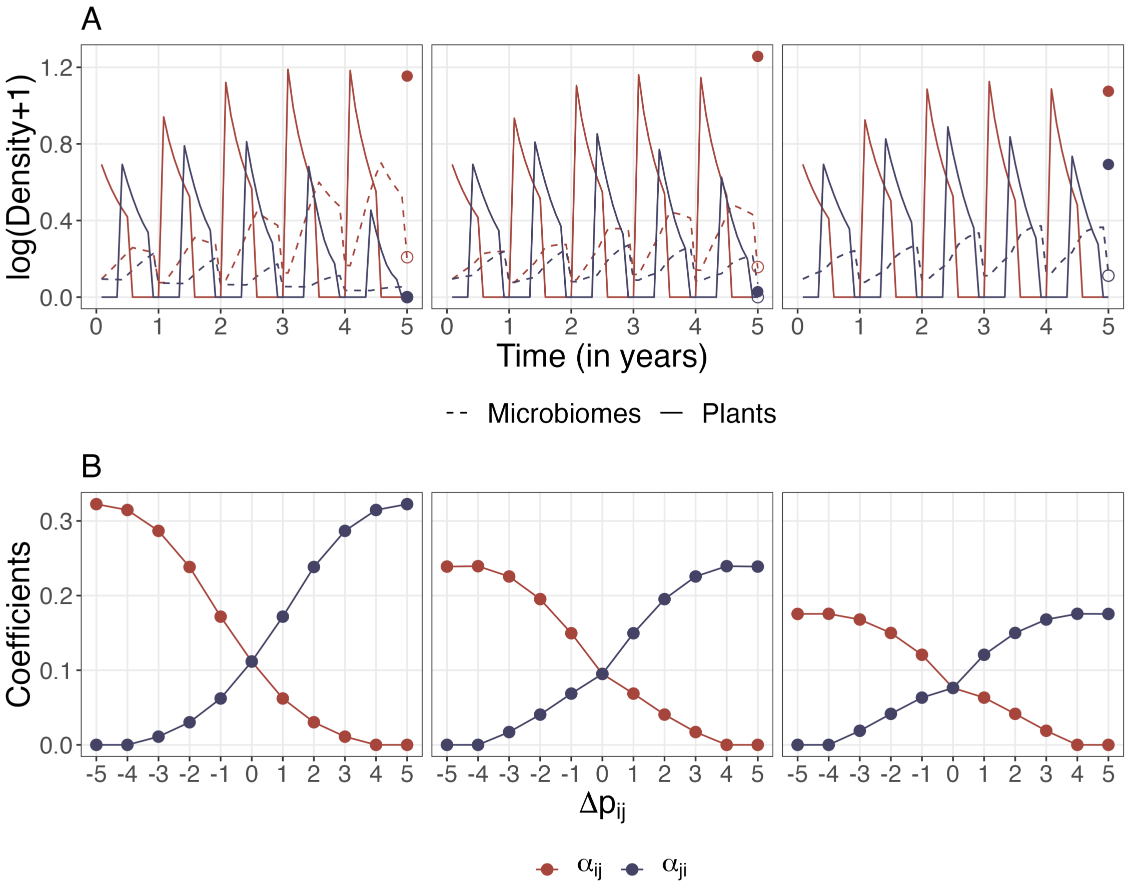

Figure S5: Two-plant communities with lower intraspecific microbial effects ($|m_{xx}| < |m_{yx}|$). From left to right: no, medium, and high overlap in associated microbiomes. A. Population dynamics of two-plant communities. Equilibrium densities (after 50 years) of plants (in seeds) and microbiomes at the end of the year are shown in solid and empty dots. B. Fitted interspecific interaction coefficients.

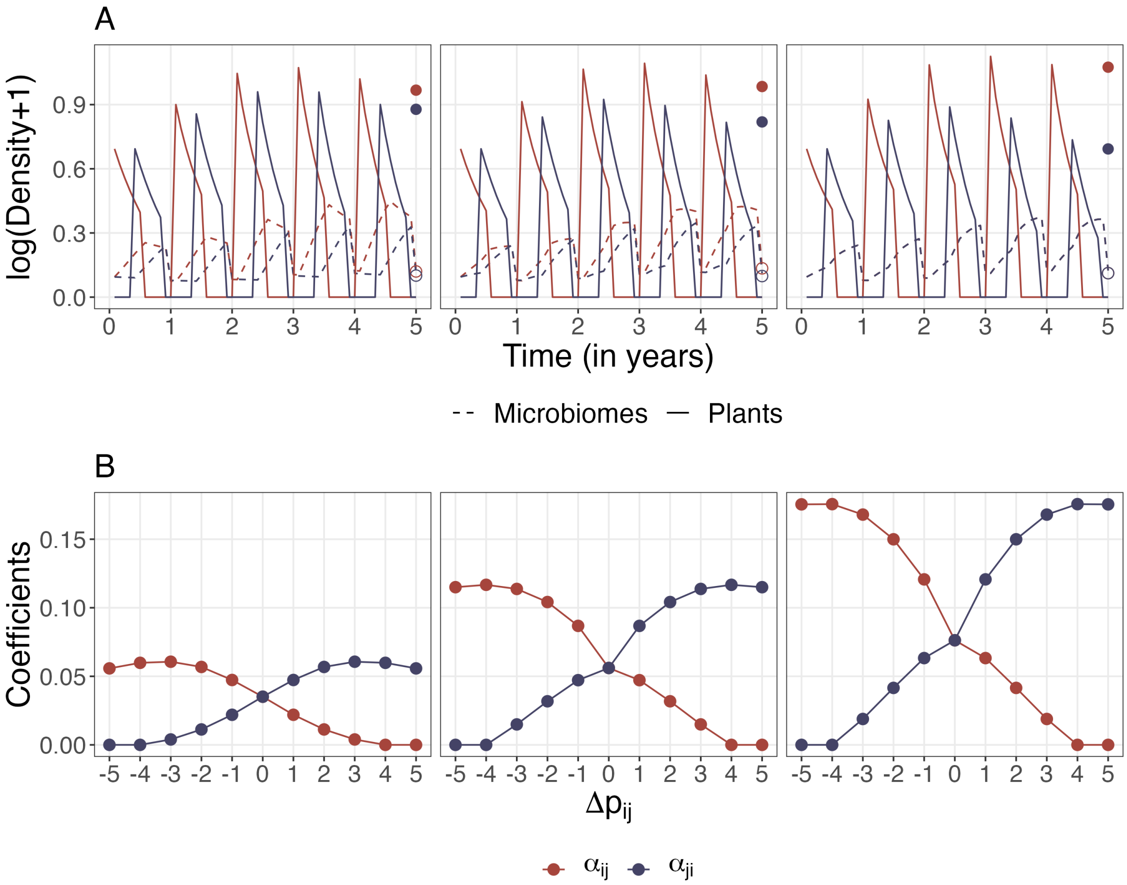

Figure S6: Two-plant communities with higher intraspecific microbial effects ($|m_{xx}| > |m_{yx}|$). From left to right: no, medium, and high overlap in associated microbiomes. A. Population dynamics of two-plant communities. Equilibrium densities (after 50 years) of plants (in seeds) and microbiomes at the end of the year are shown in solid and empty dots. B. Fitted interspecific interaction coefficients.

*Three-plant Community*

Dynamics of three-plant communities share much similarities and mechanisms of two-plant communities: when one plant's microbiome specializes on other plants $|m_{xx}| < |m_{yx}|$, increasing overlap promotes coexistence (Figure S7), and when this relationship is reversed higher overlap promotes monodominance (Figure S8).

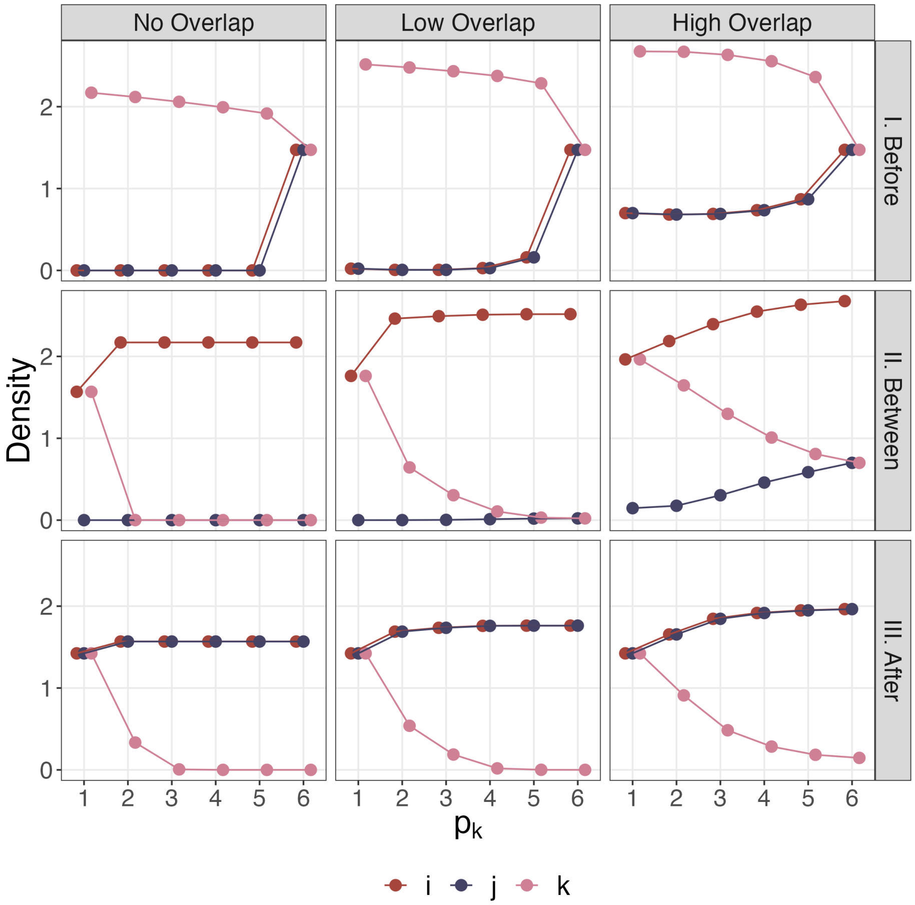

Figure S7: Three-plant communities with lower intraspecific microbial effects ($|m_{xx}| < |m_{yx}|$). Equilibrium population (after 50 years) of three plant species in three-plant communities by cases. Scenario I shows plant $k$ arriving before the focal pair i and j, Scenario II shows $k$ arriving between the focal pair, and Scenario III shows $k$ arriving after the focal pair.

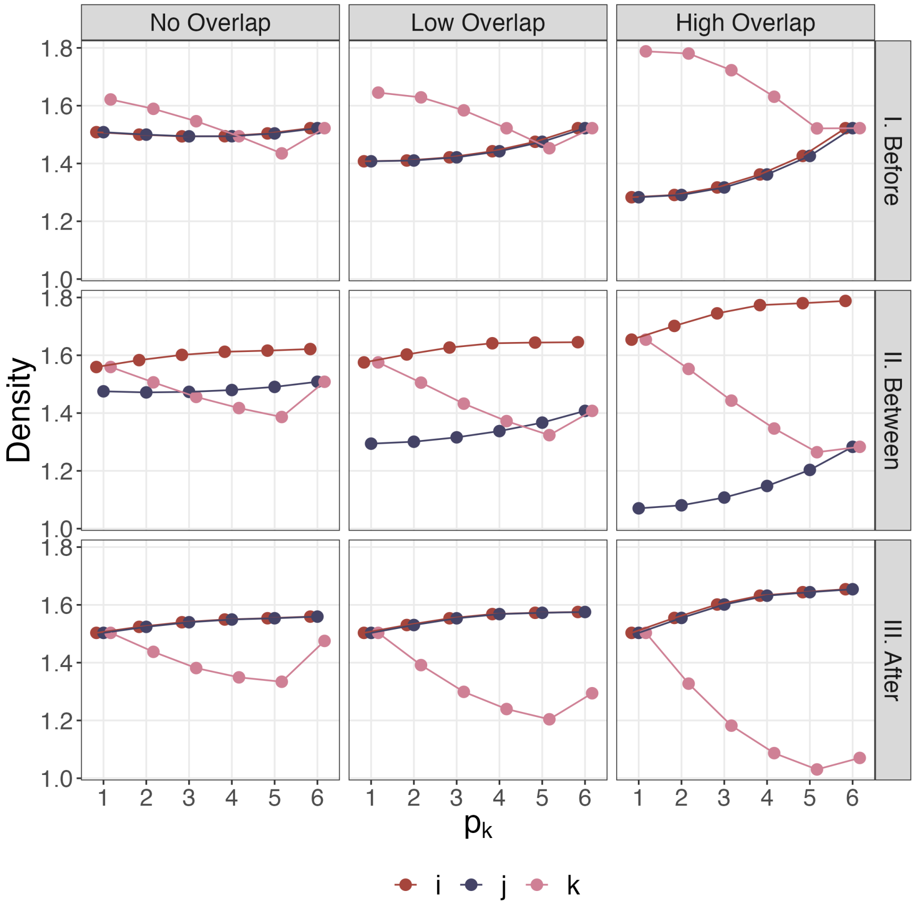

Figure S8: Three-plant communities with higher intraspecific microbial effects ($|m_{xx}| > |m_{yx}|$). Equilibrium population (after 50 years) of three plant species in three-plant communities by cases. Scenario I shows plant $k$ arriving before the focal pair i and j, Scenario II shows $k$ arriving between the focal pair, and Scenario III shows $k$ arriving after the focal pair.

Although overlap in microbiomes is linked to both coexistence and the magnitude of higher-order interactions, patterns of time-dependent interaction modifications are qualitatively the same, and no clear link can be drawn between their magnitudes and coexistence (Figure S9-S10).

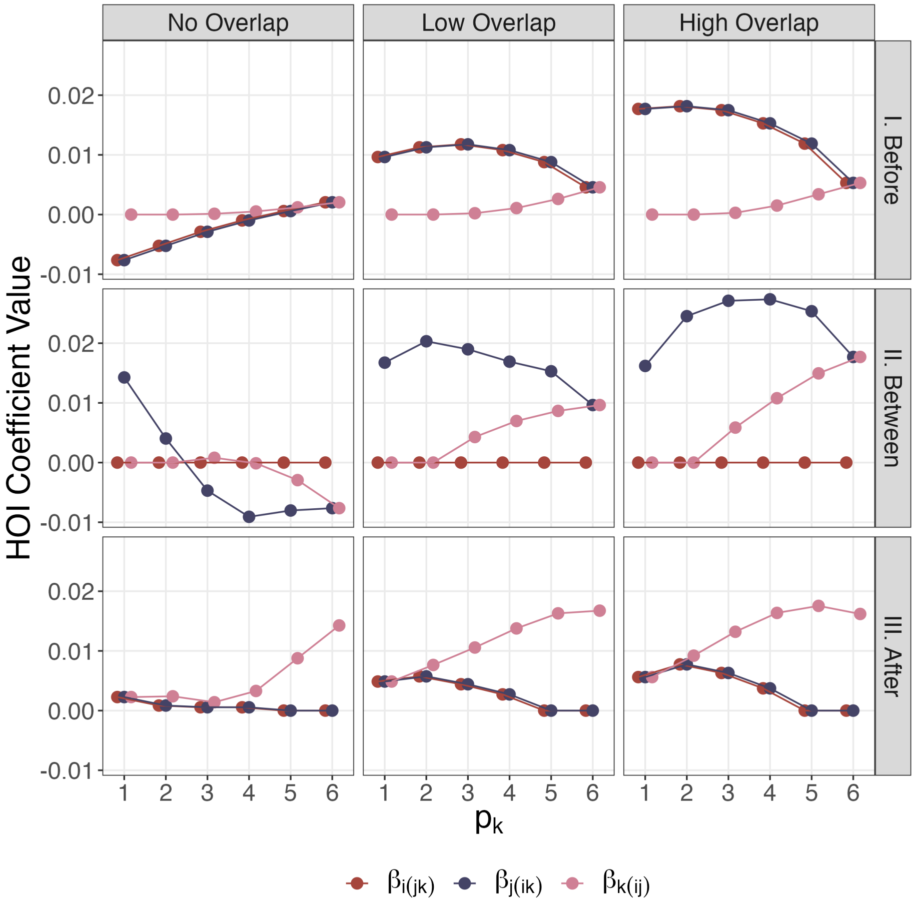

Figure S9: Time-dependent interaction modification in three-plant communities with lower intraspecific microbial effects ($|m_{xx}| < |m_{yx}|$) and different levels of overlap. The higher-order interaction coefficients $\beta_{i(jk)}$, $\beta_{j(ik)}$, and $\beta_{k(ij)}$ are fitted according to Eqn. 8.

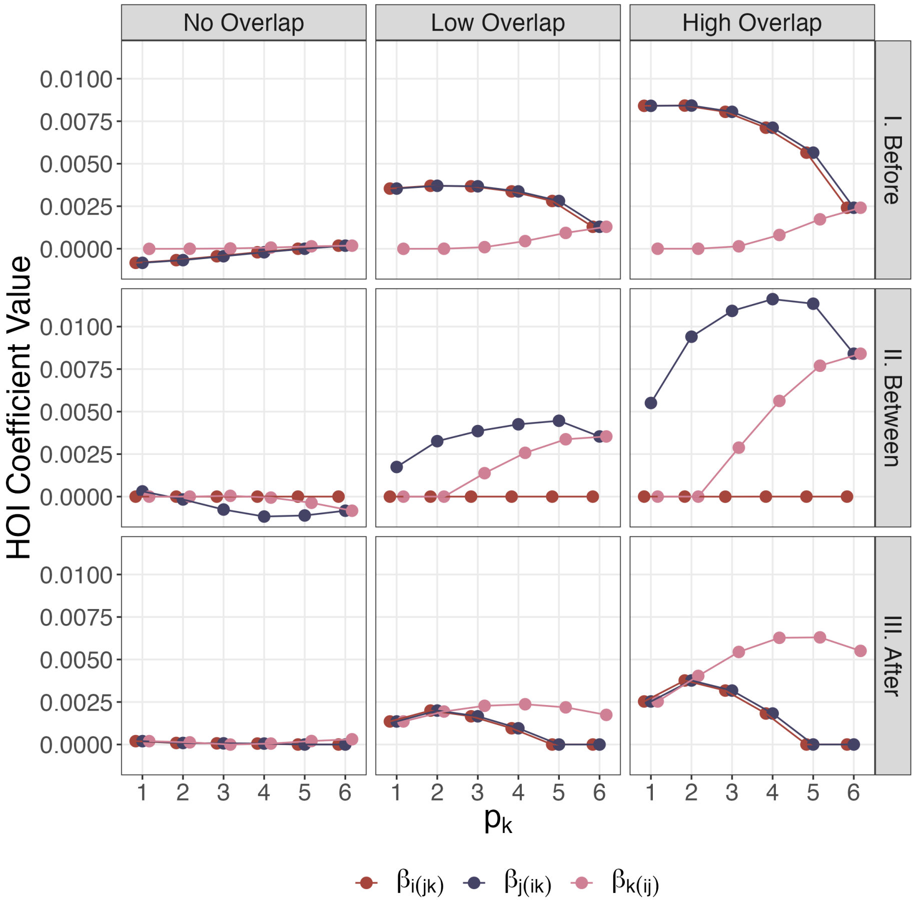

Figure S10: Time-dependent interaction modification in three-plant communities with higher intraspecific microbial effects ($|m_{xx}| > |m_{yx}|$) and different levels of overlap. The higher-order interaction coefficients $\beta_{i(jk)}$, $\beta_{j(ik)}$, and $\beta_{k(ij)}$ are fitted according to Eqn. 8.

#### *Other Higher-order Coefficients*

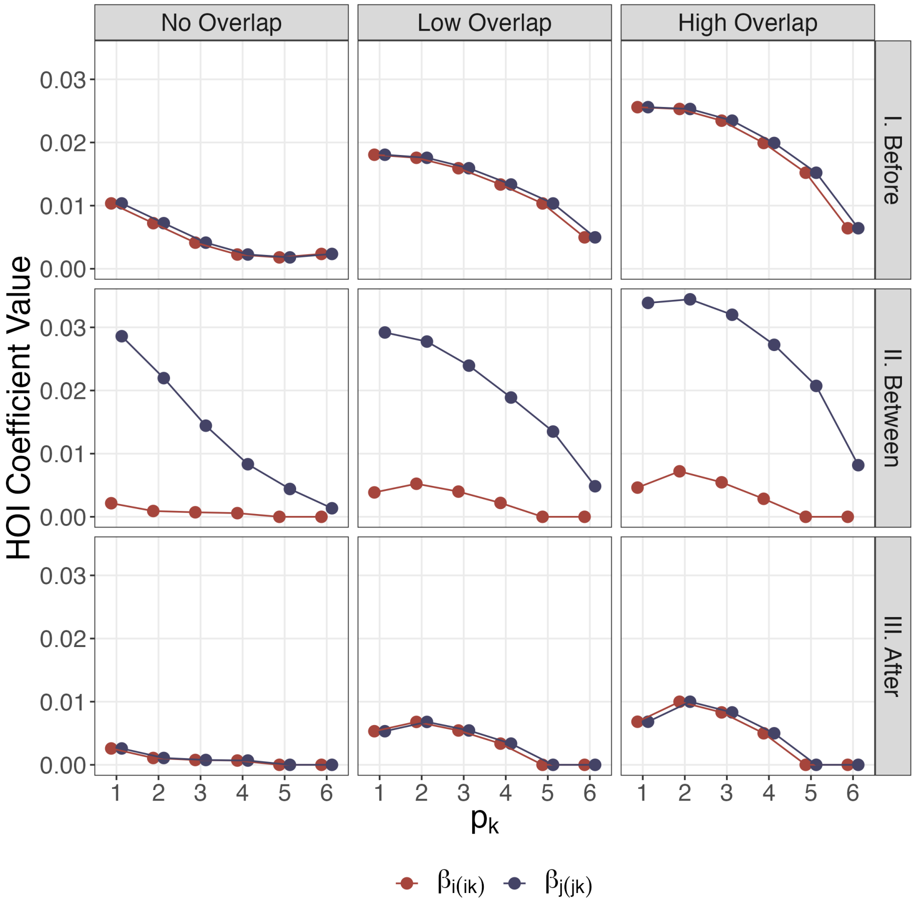

Figure S11: Time-dependent interaction modification in three-plant communities by cases. The higher-order interaction coefficients $\beta_{i(ik)}$ and $\beta_{j(jk)}$ are fitted according to Eqn. 8.

#### *Random Parameters*

Here, we test the robustness of our results by first randomly drawing 100 sets of total capacity of cultivation for each plant $V_x$ and microbial effects to plants $m_{xx}$ from two uniform distributions. In rare cases that the randomly drawn parameters produce unbounded population dynamics at the first year. They are omitted in the model fitting process. In our trials this is usually less than 2% of the simulations. We do not observe any qualitative changes in patterns of time-dependent interaction modifications (Figure S13-S14).

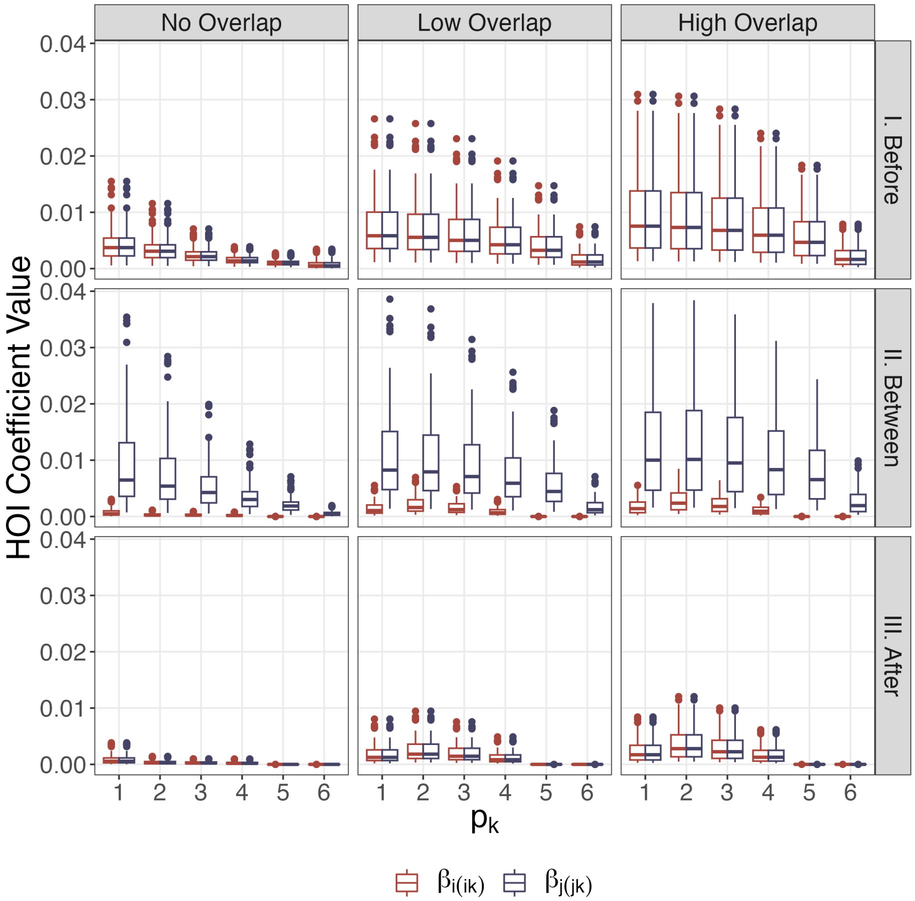

Figure S12: Time-dependent interaction modification in three-plant communities by cases, fitted from population dynamics generated by 100 random selections of total cultivation rates ($V$) and microbial effects ($m$). The higher-order interaction coefficient $\beta_{i(ik)}$ and $\beta_{j(jk)}$ are fitted according to Eqn. 8.

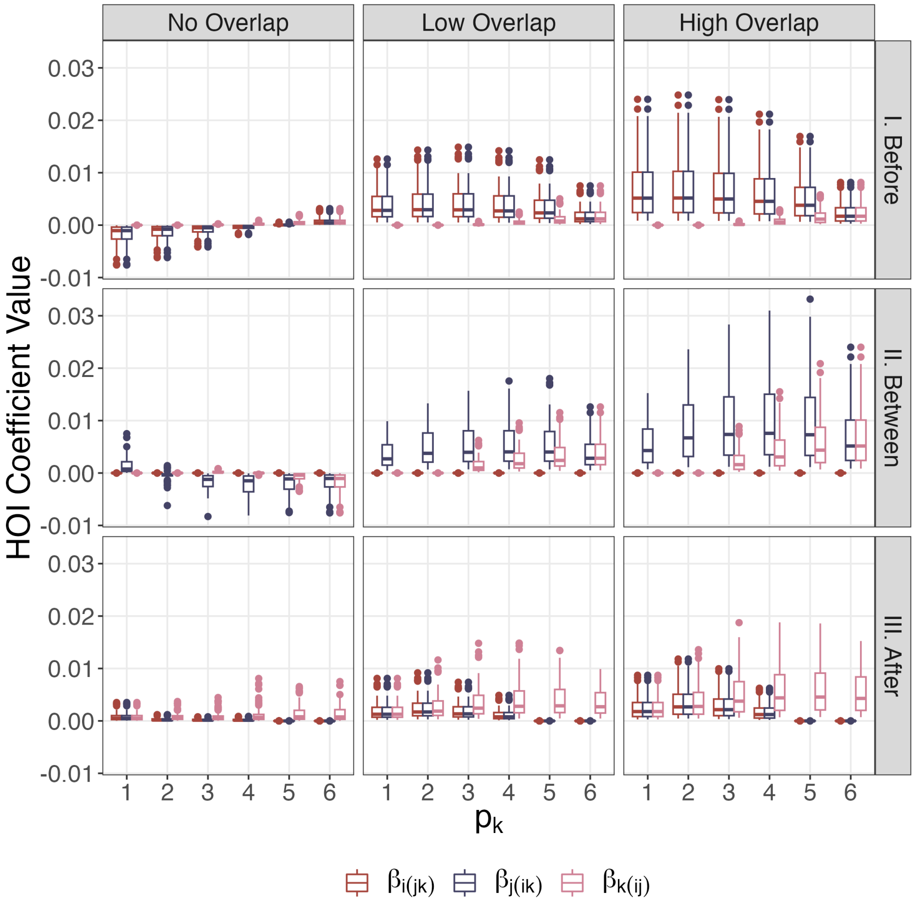

Figure S13: Time-dependent interaction modification in three-plant communities by cases, fitted from population dynamics generated by 100 random selections of total cultivation rates ($V$) and microbial effects ($m$). The higher-order interaction coefficient $\beta_{i(jk)}$, $\beta_{j(ik)}$, and $\beta_{k(ij)}$ are fitted according to Eqn. 8.

We then use a metaanalysis of plant-soil feedback experiments (Yan et al. 2022) to inform the relationship between the microbial effect from a microbiome to its own host, $m_{xx}$, and to other plants in the community, $m_{yx}$. We first filter all species pairs from experiments using sterile soil as reference with negative $m_{xx}$ and $m_{yx}$ values since we focus on negative feedback from microbiomes. We then calculate the ratio of intra- vs. interspecific feedback for each pair, $r = m_{xx}/m_{yx}$. From the data, $r \sim lognormal(0.109, 1.168)$, meaning that $m_{xx}$ tends to be stronger (more negative) than $m_{yx}$. To incorporate this data into our model, we fix the total capacity of effects from each microbiome (similar to fixing each plant's ability of cultivating microbiomes): $M_x = m_{xx} + \sum_y m_{yx}$ for all $y \neq x$. We set $M_x = -0.9$ because in the standard scenario, all intra- and interspecific microbial effects are set to $-0.3$. We first generate proportion $r$ from the above lognormal distribution, then partition $M_x$ according to $r$. Interspecific microbial effects are equally split between plants. e.g., in a three-plant community, for microbiome associated with plant $i$, $M_i = m_{ii} + rm_{ii}$, and $m_{ji} = m_{ki} = rm_{ii}/2$. We repeat this process for 100 times. In rare cases that the randomly drawn parameters produce unbounded population dynamics at the first year. They are omitted in the model fitting process. In our trials this is usually less than 2% of the simulations. We do not observe any qualitative changes in patterns of time-dependent interaction modifications (Figure S15-S16).

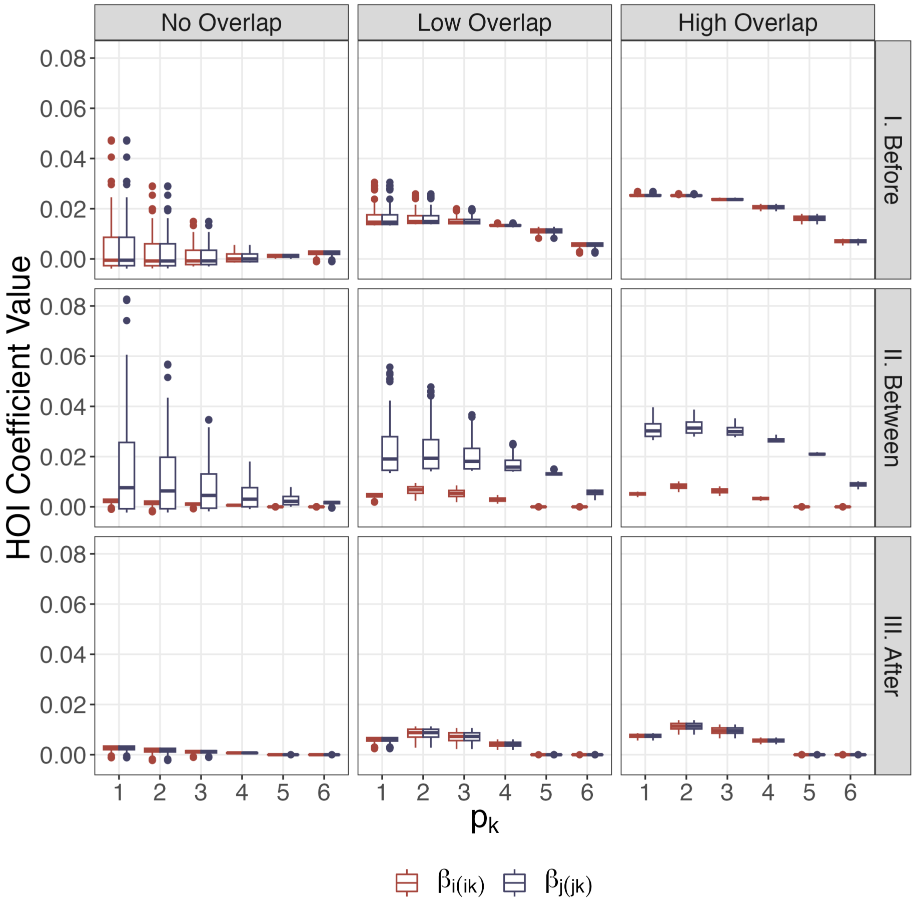

Figure S14: Time-dependent interaction modification in three-plant communities by cases, fitted from population dynamics generated by 100 random selections of microbial effects ($m$) from the distribution of a metaanalysis of empirical studies. The higher-order interaction coefficient $\beta_{i(ik)}$ and $\beta_{j(jk)}$ are fitted according to Eqn. 8.

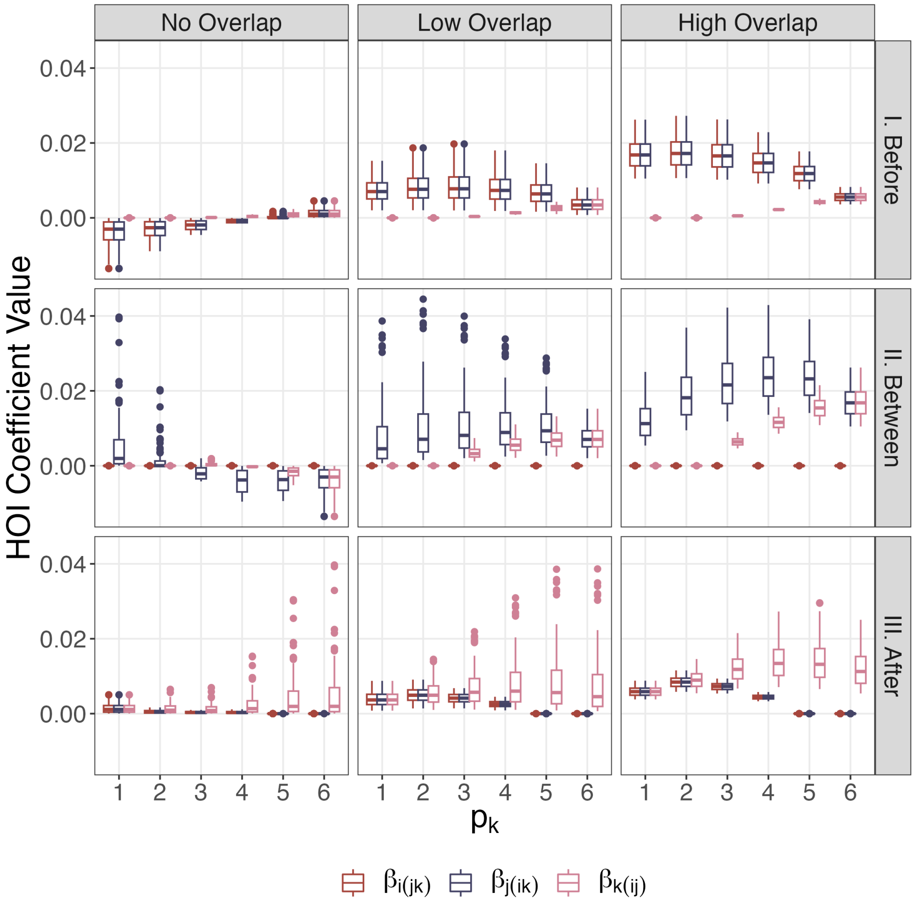

Figure S15: Time-dependent interaction modification in three-plant communities by cases, fitted from population dynamics generated by 100 random selections of microbial effects ($m$) from the distribution of a metaanalysis of empirical studies. The higher-order interaction coefficient $\beta_{i(jk)}$, $\beta_{j(ik)}$, and $\beta_{k(ij)}$ are fitted according to Eqn. 8.

#### *Model Performance*

To evaluate the performance of the model with higher-order interaction terms (Eqn. 8), we calculated the root-mean-squared errors (RMSEs) and Akaike Information Criterion (AIC) of pairwise vs. higher-order models using either plant $i$ or $j$ as the focal species. We visualize the performance as the difference between pairwise and higher-order models such that positive values indicate better performance. Comparing RMSEs, the models containing higher-order interactions terms almost always fits better than pairwise models (Figure S17). The performance improvement is larger with higher overlap in microbiomes, and its relationship with the phenology of plant $k$ loosely resembles that observed with the fitted coefficients of higher-order interactions: differences in model performance are highest with largest fitted higher-order coefficients and often approaches 0 when fitted higher-order coefficients are close to 0 (Figure 4). Differences of AIC values between pairwise and higher-order models are qualitatively identical to patterns of RMSE (Figure S18). When fitted higher-order coefficients are close to 0, the pairwise model has lower AIC, indicating that the two models perform similarly, and the higher-order model is penalized for more parameters. These results reiterate our findings that strong time-dependent interaction modifications occur with increasing level of overlap in microbiomes.

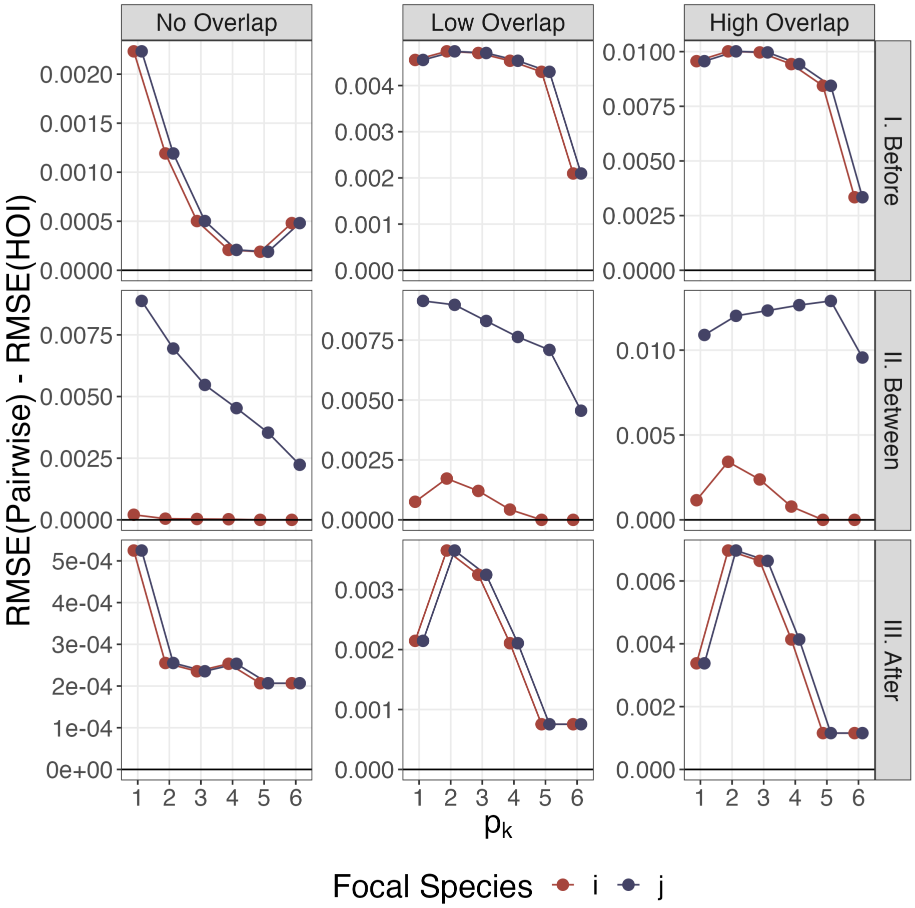

Figure S16: Difference of root-mean-squared errors (RMSE) between pairwise (Eqn. 5) and higher-order (Eqn. 8) Ricker models when fitted to population dynamics of three-plant communities. A value above 0 means the higher-order model performs better.

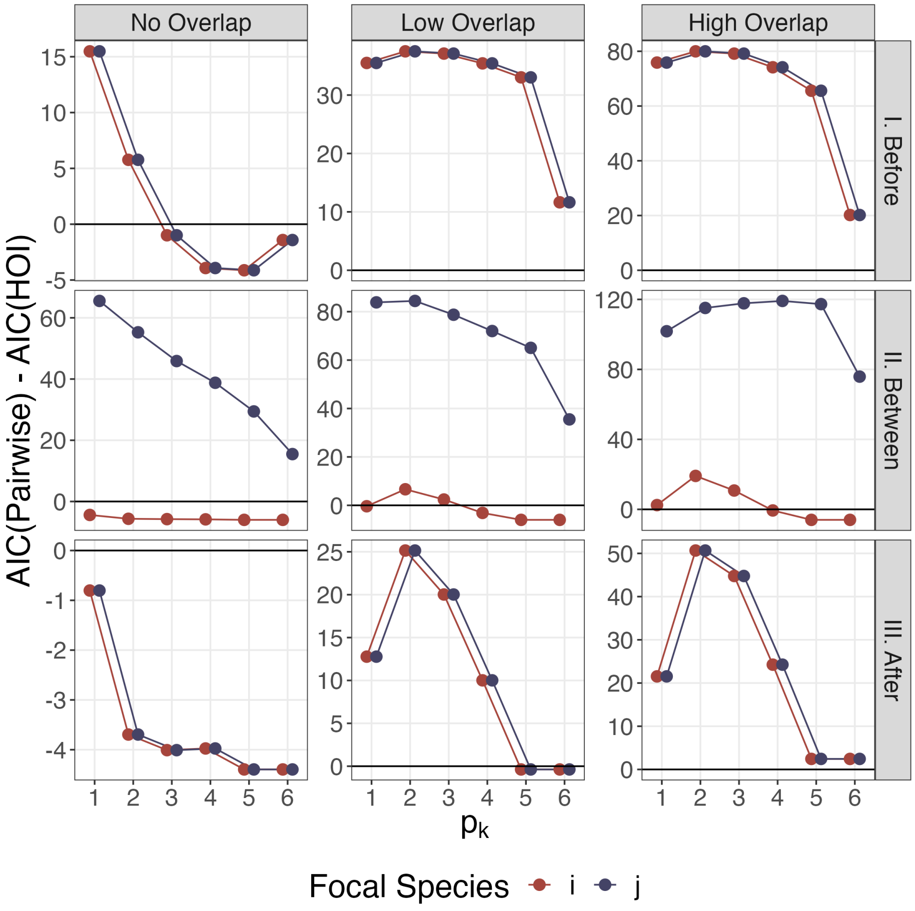

Figure S17: Difference of Akaike Information Criterion (AIC) between pairwise (Eqn. 5) and higher-order (Eqn. 8) Ricker models when fitted to population dynamics of three-plant communities. A value above 0 means the higher-order model performs better.

### Appendix II: Alternative Method of Calculating Time-dependent Interaction Modifications

#### *The Difference Method*

Consider a system with three competing species following a Ricker model:

$$\frac{N_i(t+1)}{N_i(t)} = \lambda_i \exp(-\sum \alpha_{ij} N_j(t)), \tag{S.5}$$

where $\lambda_i$ is the intrinsic growth rate of species $i$, and $\alpha_{ij}$ is the per-capita interaction from species $j$ to $i$.

In a pairwise system with trait-dependent priority effects, the per-capita interaction from species $j$ to $i$ is dependent on the difference between their arrival times within a year (phenology), $p_i$ and $p_j$, which can be further considered as the sum of a baseline interaction $\alpha_{ij|0}$ and a change in interaction strength that depends on the difference between $p_i$ and $p_j$:

$$\alpha_{ij} = \alpha_{ij|0} + g_{ij}(p_j - p_i). \tag{S.6}$$

We define this phenological difference $p_j - p_i$ as $\Delta p_{ij}$. Note that this definition is equivalent to Eqn. 6 in the main text but just decomposes the total per-capita interspecific interaction, $\alpha_{ij}$, into a constant and time-dependent term ($g_{ij}(\Delta p_{ij})$).

With time-dependent interaction modification, *both* the presence and the timing of the third species, $p_k$ may affect the interaction between species $i$ and $j$ in addition to $\Delta p_{ij}$. Similarly, this per-capita interaction can also be written as the sum of a baseline interaction $\alpha_{ij|0}$, a function that changes with the phenological difference between species $i$ and $j$, and the phenology of $k$:

$$\alpha'_{ij} = \alpha_{ij|0} + g_{ij}(\Delta p_{ij}) + g'_k(p_k). \tag{S.7}$$

To detect time-dependent interaction modification, we examine whether the difference between pairwise interaction coefficients $\alpha_{ij}$ and $\alpha'_{ij}$:

$$\Delta\alpha_{ij} = \alpha'_{ij} - \alpha_{ij} = g'(p_k). \tag{S.8}$$

If this difference changes the phenology of species $k$ ($p_k$), then the interaction modification is also time-dependent.

#### *Connecting the Two Methods*

Here, we show that the "difference" method outlined above is mathematically connected with the method of quantifying time-dependent interaction modifications in the main text. First, we adopt the definitions of Kleinhesselink et al. (2022) and divide potential higher-order interactions in a three-species system into "soft" and "hard" higher-order interactions. The former arises when the interaction between the two species can be modified by the density of one of the species in the pair. The latter arises when the interaction between the two species can be modified by the density of a third species. For the purpose of connecting the two methods, we need to first clarify the working definitions of "*density*-dependence" and "*time*-dependence". The former refers to how densities of species affect the pairwise interaction. For instance, the term $\beta_{i(jk)}N_jN_k$ in Eqn. 7 of the main text is a form of density dependence, where the overall interaction experienced by species $i$ is dependent on the densities of species $j$ and $k$, scaled by the coefficient $\beta_{i(jk)}$. On the other hand, time-dependence refers to how the interaction coefficient, $\beta_{i(jk)}$, depends on the arrival times of species $i$, $j$, and $k$. In other words, we consider the temporal changes of species density, i.e., how $N_j$ or $N_k$ change with time, as density-dependence, but not time-dependence, because the per-capita coefficient of density dependence, $\beta_{i(jk)}$, does not change with time. This dichotomy is similar to the one between frequency-dependent and trait-dependent priority effects (Zou and Rudolf 2023).

##### *Time-dependent Modification of Pairwise Interactions*

Consider a three-species system in which the interaction between two species, $i$ and $j$, can be modified by both the density and the timing of one of the species, i.e., the overall pairwise interaction $\alpha_{ij}$ is both time-dependent and density-dependent; the latter fits the definition of a "soft" higher-order interaction.

We first write the model that displays trait-dependent priority effects. As outlined in the main text, in a Ricker model with such interaction:

$$\frac{N_i(t+1)}{N_i(t)} = \lambda_i \exp(-\alpha_{ii}N_i(t) - \alpha_{ij|0}N_j(t) - \alpha_{ik}N_k(t) - g_{ij}(\Delta p_{ij})N_j(t)). \tag{S.9}$$

Here, we follow the definition above and specify that $\alpha_{ij|0}$ is the baseline interaction coefficients between species $i$ and $j$ modified by neither time nor density of species $i$ or $j$. For simplicity of presentation, we do not similarly decompose $\alpha_{ik}$ as in Eqn. S.6.

We then write the model with a "soft" higher-order interaction between species $i$ and $j$ in the same Ricker model:

$$\frac{N_i(t+1)}{N_i(t)} = \lambda_i \exp(-\alpha_{ii}N_i(t) - \alpha_{ij|0}N_j(t) - \alpha_{ik}N_k(t) - \beta_{i(ij)}N_i(t)N_j(t)). \tag{S.10}$$

Here, the term $\beta_{i(ij)}$ is the interaction coefficient by which the density of both species $i$ and $j$ affect the overall interaction experienced by $i$.

Because we use Eqn. S.9 and Eqn. S.10 to represent the same system that exhibits both time- and density-dependent interaction modifications (i.e., "soft" higher-order interaction),

$$g_{ij}(\Delta p_{ij}) = \beta_{i(ij)}N_i(t). \tag{S.11}$$

This means that in a system that displays trait-dependent priority effects, the pairwise interaction between species $i$ and $j$ can be considered as both time-dependent and density-dependent interaction modifications. In other words, trait-dependent priority effects can be density-dependent, and "soft" higher-order interactions can be time-dependent.

*Time-dependent Modification of Higher-order Interactions*

Consider a three-species system in which the interaction between two species, $i$ and $j$, can be modified by both the density and the timing of the third species $k$, i.e., the overall pairwise interaction $\alpha_{ij}$ is both time-dependent and density-dependent; the latter fits the definition of a "hard" higher-order interaction.

We first derive the model using the "difference" method:

$$\frac{N_i(t+1)}{N_i(t)} = \lambda_i \exp(-\alpha_{ii}N_i(t) - \alpha_{ij|0}N_j(t) - \alpha_{ik}N_k(t)) - g'(p_j, p_k)N_j(t)). \tag{S.12}$$

Again, $\alpha_{ij|0}$ is the baseline interaction coefficients between species $i$ and $j$ modified by neither time nor density of species $i$ or $j$, and we do not decompose $\alpha_{ik}$ for simplicity of presentation. Following the main text, we further simplify $g'(p_j, p_k)$ to $g'(p_k)$ by assuming $p_j$ as a constant.

We then derive the model with a "hard" higher-order interaction between species $i$ and $j$ in the same Ricker model:

$$\frac{N_i(t+1)}{N_i(t)} = \lambda_i \exp(-\alpha_{ii}N_i(t) - \alpha_{ij|0}N_j(t) - \alpha_{ik}N_k(t) - \beta_{i(jk)}N_j(t)N_k(t)), \tag{S.13}$$

where $\beta_{i(jk)}N_j(t)N_k(t)$ represents the effect of both densities of $j$ and $k$ on $i$. For simplicity of presentation we do not include other higher-order interactions.

Because we use Eqn. S.12 and Eqn. S.13 to represent the same system that exhibits both time- and density-dependent interaction modifications (i.e., "hard" higher-order interactions),

$$g'(p_j, p_k) = \beta_{i(jk)}N_k(t). \tag{S.14}$$

With $p_k$ and $N_k$ on either side of the equation, this "hard" higher-order interaction term is a combination of both time-dependent and density-dependent interaction modifications.

#### *Comparing the Two Methods*

Here, we first show the calculated time-dependent interaction modifications from the "difference" method (Eqn.S8) using the same simulated data in the main text. Comparing with the fitted $\beta$ coefficients, these $\Delta\alpha$ coefficients show generally the same trend with later phenology of $k$ and larger overlap in microbiomes. However, the sign and magnitude of these time-dependent interaction modifications are different from the $\beta$ coefficients (Figure S19). To test for the robustness of these patterns, we further decreased the number of response surfaces by using three densities of plant $i$ and $j$, and one density of plant $k$, such that both two- and three-plant community has 9 density combinations (i.e., the only difference is the addition of plant $k$), while keeping the six germination phenology of plant $k$. While the general trend of interaction modifications with $p_k$ remains similar to those calculated from the "difference" method and even those fitted from a higher-order interaction model (compare Figure S20 to Figure S19 and Figure 4), the $\Delta\alpha$ values differ more in magnitude and, in some cases, show completely flipped signs.

These results illustrate the pros and cons of using the "difference" method to calculate time-dependent interaction modification. The "difference" method requires less experimental effort because a density gradient of the third species ($k$ in above sections) is not needed; one only needs to estimate the pairwise interaction between species $i$ and $j$ with the absence, and presence but at different timing of species $k$, therefore only requiring density gradients of species $i$ and $j$. Because only pairwise interaction coefficients are required, using the same model setup and parameters but only nine response surfaces (three densities of plants $i$ and $j$) we are able to recover much of the general patterns of time-dependent interaction modification in the community.

However, the remarkable differences in magnitudes and signs indicate that time-dependent interaction modifications estimated from the "difference" method are not as equivalent as $\beta$ terms fitted directly from a model with higher-order interactions, even though they are mathematically linked (see above). This is because in our plant-soil feedback model, the presence or the timing of $k$ dramatically changes community dynamics such that any pairwise model would fit poorly. Therefore, the statistical credibility of a pairwise model fitted to the focal pair in a three-plant community is low, and this impacts the credibility of calculated time-dependent interaction modifications. This can happen when other vital rates or life history traits of the focal pair are strongly affected by the third species. Furthermore, because this difference in pairwise coefficients ($\Delta\alpha_{ij}$) depends on the density of species $k$ as well, stochasticity in $N_k$ caused by natural fluctuations or experimental errors may also strongly affect the calculated time-dependent interaction modification.

Fitting a model with explicit higher-order terms (i.e., the method we illustrate in the main text) increases the credibility of time-dependent interaction modifications (but only with appropriate base model structure; see Kleinhesselink et al. 2022), but requires density gradients of all three species in addition to a temporal gradient of the third species, which is highly unfeasible. Either method can be chosen for the specific purpose of each study. We choose to directly fit a model with higher-order terms because (1) we assume a fixed total cultivation rate from plants, such that adding a third plant will decrease the rate of interspecific cultivation to each microbiome not associated with the focal plant by half and is therefore less comparable to the original two-plant community; (2) we use a simulated dataset that does not have limitations on implementing density gradients of all three species. On the other hand, despite the statistical caveats, the "difference" method may still be useful in experiments where the aim is not to precisely quantify, but determine the presence of time-dependent interaction modifications.

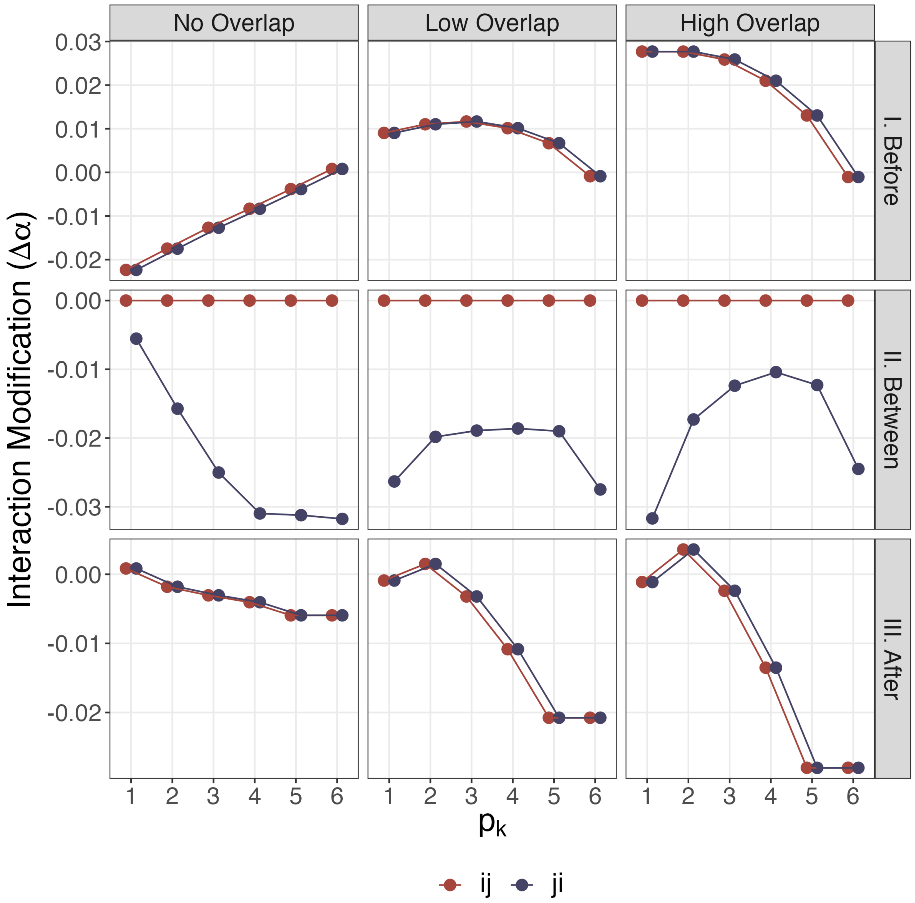

Figure S18: Time-dependent interaction modification in three-plant models by cases. The interaction modification ($\Delta\alpha$) is calculated according to Eqn. S8. Scenario I shows plant $k$ germinating before the focal pair $i$ and $j$, scenario II shows $k$ germinating between the focal pair, and scenario III shows $k$ germinating between the focal pair. When the focal pair arrives at the same time (scenarios I and II), interaction modifications on $\alpha_{ij}$ and $\alpha_{ji}$ are identical; points are jittered for visibility.

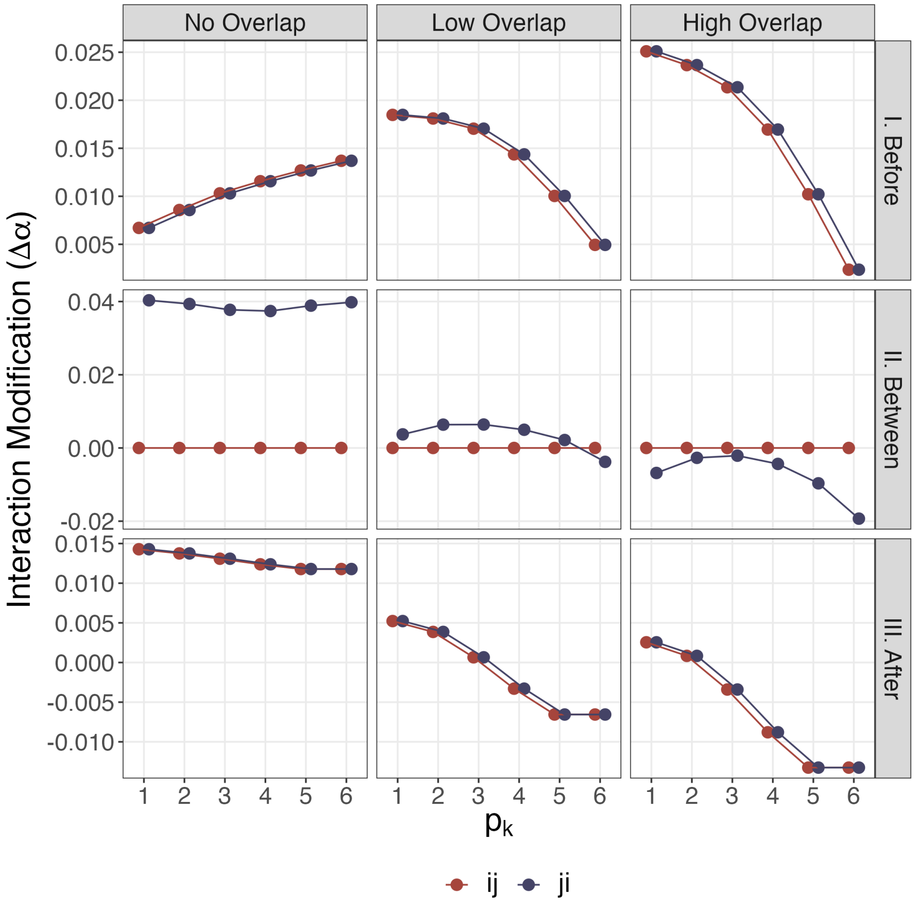

Figure S19: Time-dependent interaction modification in three-plant models by cases, with only 9 response surfaces. The interaction modification ($\Delta\alpha$) is calculated according to Eqn. S8. Scenario I shows plant $k$ germinating before the focal pair $i$ and $j$, scenario II shows $k$ germinating between the focal pair, and scenario III shows $k$ germinating between the focal pair. When the focal pair arrives at the same time (scenarios I and II), interaction modifications on $\alpha_{ij}$ and $\alpha_{ji}$ are identical; points are jittered for visibility.
